# A Comprehensive Mathematical Model of Avidity in Cytokine Signaling

**DOI:** 10.64898/2026.04.29.721617

**Authors:** Eugene F Douglass, William Bastian, Jonathan P. Mochel

## Abstract

Cytokine sensitivity varies substantially across cell types and cellular states, in part through differences in the abundance of individual receptor subunits. Yet for multicomponent receptors, the quantitative relationship between receptor abundance, binding affinity, and functional potency remains poorly defined. Here, we derive closed-form expressions for EC_50_ in multivalent ligand-receptor systems at cell surfaces. Unlike monovalent models, these equations predict that potency depends on both binding constants and receptor abundance, with individual receptor subunits playing asymmetric roles in controlling maximal complex formation and cellular sensitivity.

We validate these predictions using quantitative antibody-binding, cytokine-binding, and signaling data, and extend the equilibrium framework to steady-state signaling conditions in which downstream kinetic processes can impose additional limits on functional potency. We then derive a regression-compatible formulation and test its predicted ligand-receptor-response relationships across an *in vivo* murine cytokine perturbation atlas, human spatial transcriptomic data, and an independent cohort of 510 patients with lung adenocarcinoma. Multivariable cytokine relationships were conserved across human datasets and associated with clinical outcome. In addition, quantitative proteomics from primary human immune cells supported receptor mRNA abundance as an informative but imperfect proxy for protein abundance. Together, this framework connects multivalent receptor biophysics with cell-type-specific signaling across molecular, cellular, tissue, and patient scales.

## Introduction

### Intercellular communication

is central to normal physiology and to diseases including inflammation, fibrosis, and cancer.^1–3^ Recent advances in single-cell RNA sequencing (scRNA-seq) allow three major components of these signaling systems to be measured simultaneously: ligand expression, receptor expression, and downstream transcriptional responses.^4–6^ As illustrated in Figure 1A, ligands can be measured in “emitter cells,” while receptors and downstream signaling responses can be measured in “receiver cells.” However, dissociated single-cell data lack the spatial information needed to determine which cells are positioned to communicate. Recent studies demonstrated that cytokines typically act over length scales of approximately 20-100 μm, depending on factors including cytokine concentration at the emitter source ([C]), and cytokine sensitivity (EC_50_) of the receiver cells.^7, 8^

**Figure 1.**
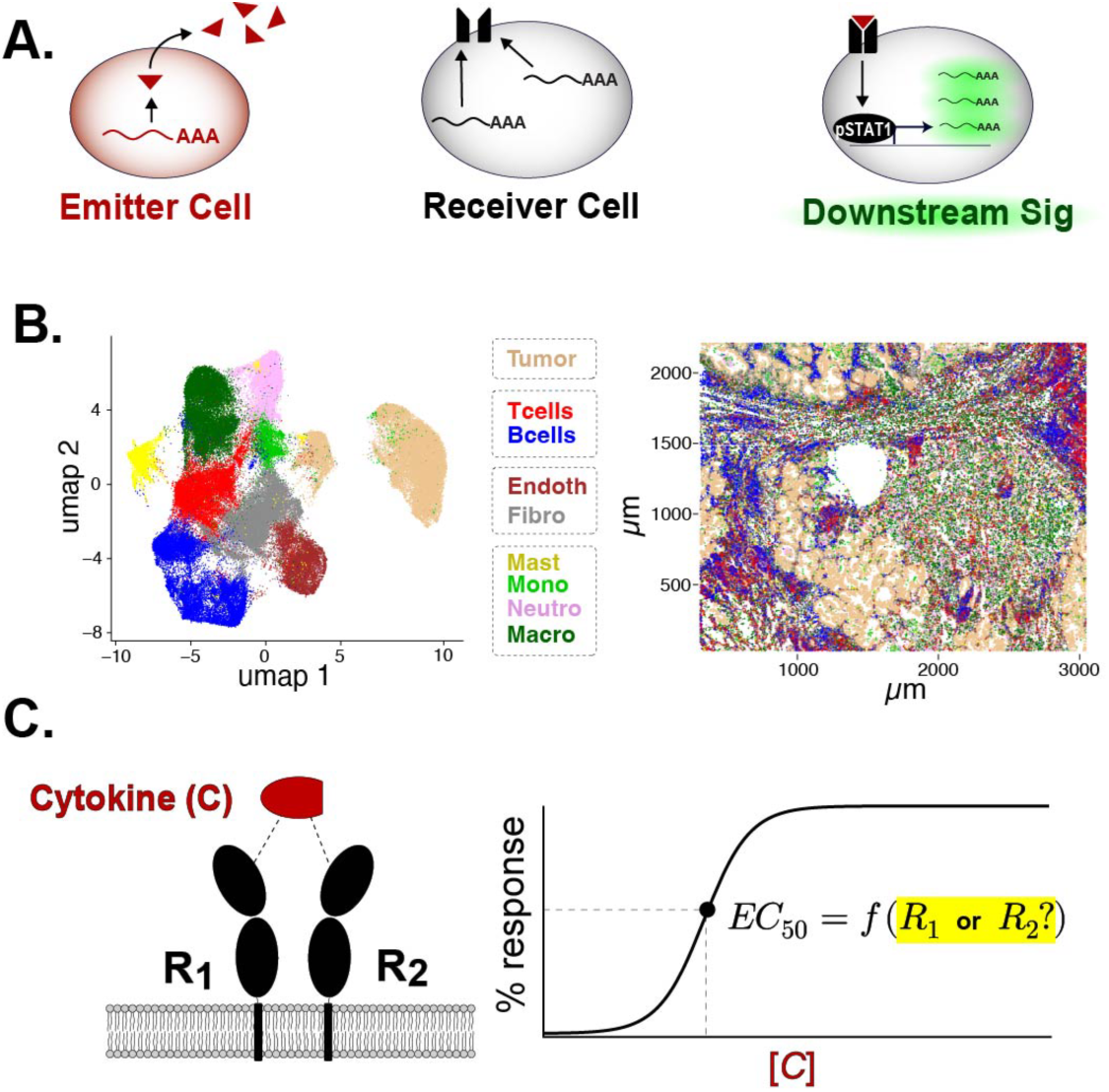
Transcriptomic evidence for modeling cytokine signaling across spatial and clinical datasets. **(A)** Single-cell transcriptomics provides three complementary measurements of cytokine signaling: ligand expression, expression of multiple receptor subunits, and downstream transcriptional activity. **(B)** CosMx Spatial Molecular Imaging simultaneously resolves these measurements together with cell identity and two-dimensional tissue location. UMAP visualization captures transcriptional cell states, whereas the in situ map preserves the physical distances between cells, an important constraint on local cytokine signaling. **(C)** The avidity framework was designed to integrate these signaling measurements across both single-cell spatial transcriptomic datasets and large bulk RNAseq cohorst. Sc-spatial data enables physically constrained analysis of ligand-receptor interactions. Bulk RNAseq cohorts, enable assessment of clinical relevance. The framework operates directly on ligand, receptor, and downstream signaling measurements. without requiring predefined sender–receiver cell-type assignments.

### Spatial Transcriptomics

Single-cell spatial transcriptomics provides this missing spatial context.^9^ In 2022, CosMx Spatial Molecular Imaging (SMI) enabled simultaneous measurement of 1,000 mRNAs and the locations of 100,416 individual cells in a human lung adenocarcinoma specimen (Fig 1B).^10^ These measurements revealed recurrent “spatial niches” defined by local cellular architecture, including tumor-cell niches, a T-cell/B-cell niche, fibroblast/endothelial regions, and a myeloid-enriched niche. Within such niches, spatial data can identify which cells are close enough to interact and quantify local cytokine expression.

However, proximity and ligand abundance alone do not determine signaling. A second question remains: *<u>How sensitive are different cells to a given cytokine</u> <u>within the same niche?</u>* Cellular sensitivity is conventionally quantified by the EC_50_, the effective cytokine concentration required to produce a 50% response,^11^ and is typically measured using ex vivo dose-response experiments fit with a Hill-type equation:^12–16^

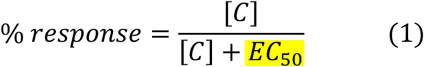

### Avidity Conceptual Challenge

For multicomponent cytokine receptors (Fig 1C), EC_50_ is not simply a fixed property of the cytokine. Many cell-cell communication methods nevertheless treat EC_50_ as constant across cytokines and cell types. While this approximation can be appropriate for simple 1:1 ligand-receptor interactions, most cytokines engage multiple receptor subunits (R_1_, R_2_),^3, 17–19^ and cellular sensitivity can depend strongly on their relative abundance (Fig 1C).^12, 20–25^ For example, approximately 100-fold differences in IL2RA (CD25) expression have been associated with approximately 100-fold shifts in the functional EC_50_ of T cells to IL-2.^13–16, 26^ Similar receptor-dependent shifts have been reported across major cytokine systems,^13, 25, 27–31^ and a recent review has emphasized that a single receptor typically dominate cell-type-specific cytokine sensitivity.^12^ Which receptor controls sensitivity, however, differs among cytokine systems and cannot generally be inferred from receptor expression alone. ^12^ Thus, even when transcriptomic data measure the ligand, each receptor subunit, and the downstream response (Fig 1A), there is no simple quantitative rule for determining how these measurements should combine to predict signaling.

### History of “Avidity”

The distinction between intrinsic binding affinity and system-dependent functional potency (EC_50_) has been recognized for more than a century.^32^ Ehrlich introduced the term “avidity” in 1897 to describe dose-dependent toxin neutralization, initially using it interchangeably with affinity,^32^ while Krauss and Doerr later proposed that avidity represented a biologically distinct property (1905).^33^ Quantitative studies in the 1950s strengthened this distinction by showing >100-fold enhancement of antibody-antigen binding at cell surfaces relative to solution.^34–36^ These observations ultimately motivated the distinction between “intrinsic affinity” (K_a_ or 1/K_d_) for monovalent interactions and “functional affinity” or “avidity” (EC_50_) for multivalent binding.^37, 38^ Despite this long history, avidity remains inconsistently defined. Several quantitative frameworks have described multivalent binding at biological surfaces and synthetic interfaces,^39^ including models incorporating receptor abundance and heterogeneous binding components.^40^

### Current Quantitative Frameworks

Most quantitative receptor models remain rooted in Clark’s receptor-occupancy framework, which describes a 1:1 interaction between a ligand (L) and a single receptor (R) using the equilibrium dissociation constant, K_d_.^41^ Under this model, the ligand concentration producing half-maximal receptor occupancy is equal to K_d_: ^42, 43^

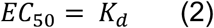

The Hill equation generalizes this framework to sigmoidal dose-response relationships and can incorporate cooperativity, but treats EC_50_ as an empirical fitting parameter rather than deriving it mechanistically.^44, 45^ Both formulations assume monovalent binding. However, most ligands at cell surfaces engage multiple receptor subunits simultaneously (Fig 1C).^15,16^ Both approaches are therefore most natural for monovalent interactions. At cell surfaces, however, most cytokines engage multiple receptor subunits simultaneously.^3, 17–19^ In these multivalent systems, EC_50_ is no longer determined by binding constants alone but can also depend on receptor abundance, which varies across cell types, activation states, and disease contexts.^46–48^ A quantitative framework that explicitly derives this receptor-density dependence is therefore needed to connect molecular binding properties with cell-type-specific sensitivity.^25, 49, 50^

### Work Presented Here

Here, we derive and validate an avidity-based framework for multicomponent cytokine signaling at cell surfaces. Building on analytical models of multicomponent binding,^51, 52^ we derive closed-form expressions that relate EC_50_ to individual binding constants and receptor abundance, answering the question posed in Figure 1C:

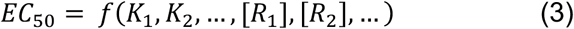

These equations predict that receptor subunits can play asymmetric roles: only one contributes to cell-type sensitivity (equation 3) while the other dictates the maximum ligand-receptor complex that can form. We first test these predictions using antibody-binding and cytokine-signaling datasets in which affinity, receptor abundance, and functional EC_50_ have been experimentally measured (Fig 2-3). We then extend the equilibrium framework to steady-state signaling, allowing downstream kinetic processes to modify functional potency while preserving the underlying ligand-receptor dependencies.

**Figure 2.**
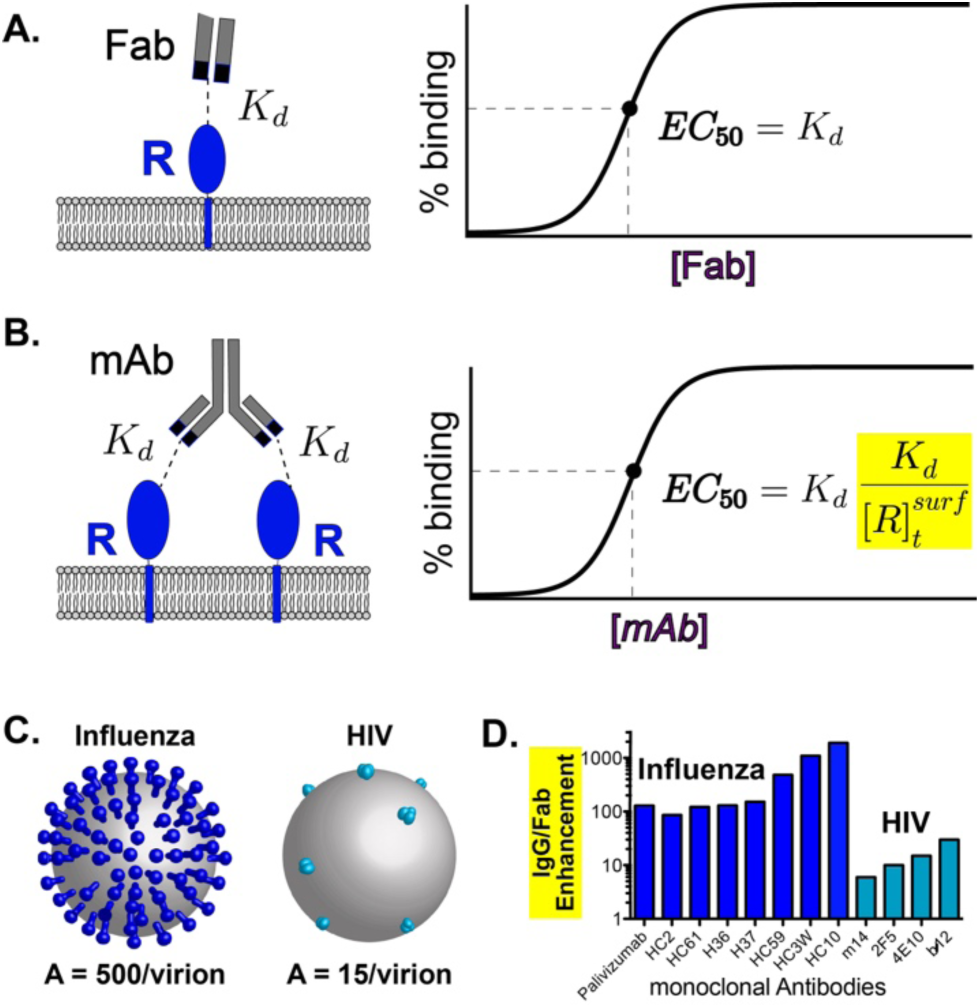
Historical and mechanistic context for antibody binding constants and avidity. **(A)** Under simple monovalent equilibrium binding, half-maximal receptor occupancy occurs at Kd; Hill-type models use EC50 as an empirical dose-response parameter **(B)** In contrast, multivalent ligands like antibodies engage multiple receptors, making functional potency (EC_50_) a composite property of receptor expression and binding avidity. **(C)** HIV spike proteins are expressed at ~10-fold lower surface density compared to other enveloped viruses, a recognized immune-evasion mechanism that limits effective antibody engagement.^53^ **(D)** Studies quantifying neutralization by full IgG versus Fab fragments have shown significantly reduced potency for monovalent Fabs, illustrating the critical role of avidity.^49, 54^ The IgG:Fab EC_50_ ratio has emerged as an empirical measure of this avidity effect.

**Figure 3.**
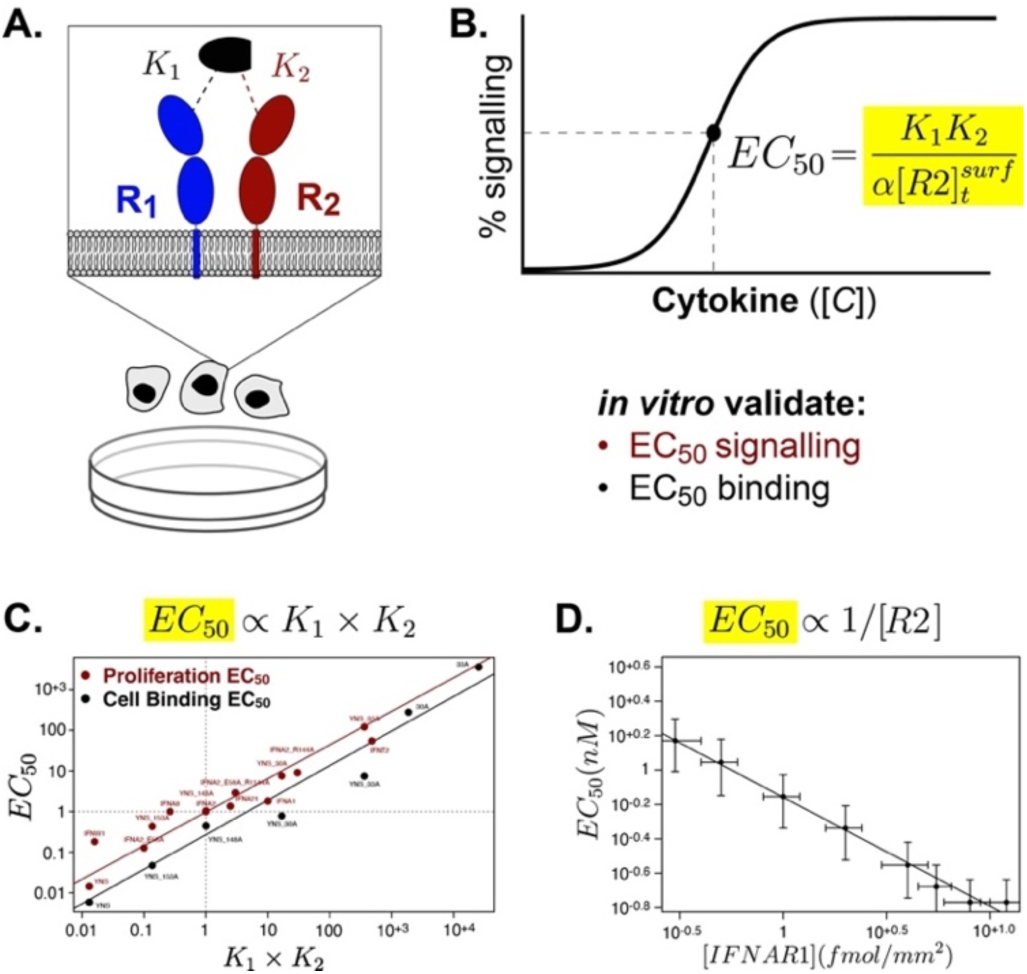
Summary of biophysical, cellular binding, and functional signaling studies on type I interferons. **(A)** Most cytokines—including type I interferons—signal through two distinct receptors (R_1_ and R_2_), each with its own binding constants (K_1_, K_2_), forming a ternary signaling complex. **(B)** Our derived EC50 equation incorporates both affinities and receptor expression to accurately model potency across diverse biological contexts. **(C)** Experimental measurements show that both cell-surface binding EC_50_ values (in black) and downstream anti-proliferative effects (in red) correlate with the product of K_1_ and K_2_, validating the model’s structure-function basis.^20, 21, 63^ **(D)** Among the two receptors, only expression of the higher-abundance “excess” receptor (R_2_) shows a significant correlation with interferon potency, while the limiting receptor (R_1_) does not—highlighting the importance of receptor balance in modulating cellular sensitivity.^23^

Finally, we translate these relationships into a regression-compatible formulation that can be tested using transcriptomic data. We evaluate the resulting predictions across an in vivo murine cytokine-perturbation atlas, human single-cell spatial transcriptomics, and an independent cohort of 510 patients with lung adenocarcinoma (Fig 4-5). We additionally use quantitative proteomics from primary human immune-cell populations to assess whether receptor mRNA abundance provides useful information about relative protein abundance. Together, these analyses test whether the receptor-dependent principles that govern cytokine potency in controlled experimental systems remain detectable across cell types, tissues, and patients.

**Figure 4.**
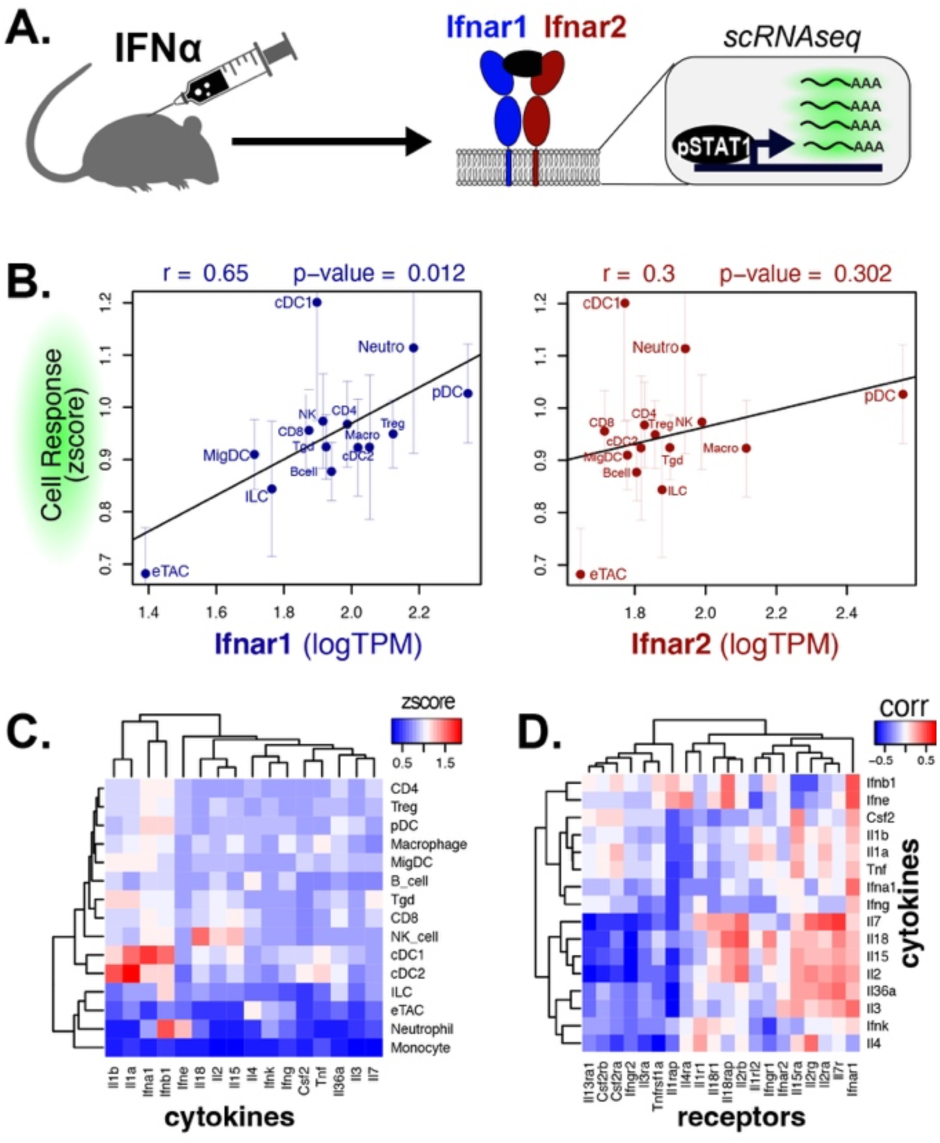
Validation of Cytokine Potency Framework Using Mouse scRNA-seq Data from *In Vivo* Cytokine Stimulation.^71^. **(A)** Schematic overview of the experimental design from Cui et al., in which 86 recombinant cytokines were injected into mice and draining lymph nodes were harvested for single-cell RNA-seq analysis. Each cell type’s baseline receptor expression and induced gene expression were quantified relative to PBS-injected controls. **(B)** Analysis of type I interferon signaling across 18 immune cell types shows a strong correlation between Ifnar1 expression and interferon-induced gene expression, while Ifnar2 shows a weaker, non-significant correlation. **(C)** Heatmap summarizing the average magnitude of gene induction for the 16 cytokines with the most consistent and robust responses across all immune cell types. **(D)** Correlation matrix comparing receptor expression (columns) with cytokine-induced gene expression (rows) across the 16 high-signal cytokines. Only 50% of receptors display positive correlations with downstream signaling. In most cytokines, one co-receptor shows a strong positive correlation, while the other is uncorrelated or inversely correlated, highlighting asymmetric control of cytokine sensitivity. Correlations in panels B-D are exploratory and were not adjusted for multiple testing.

**Figure 5.**
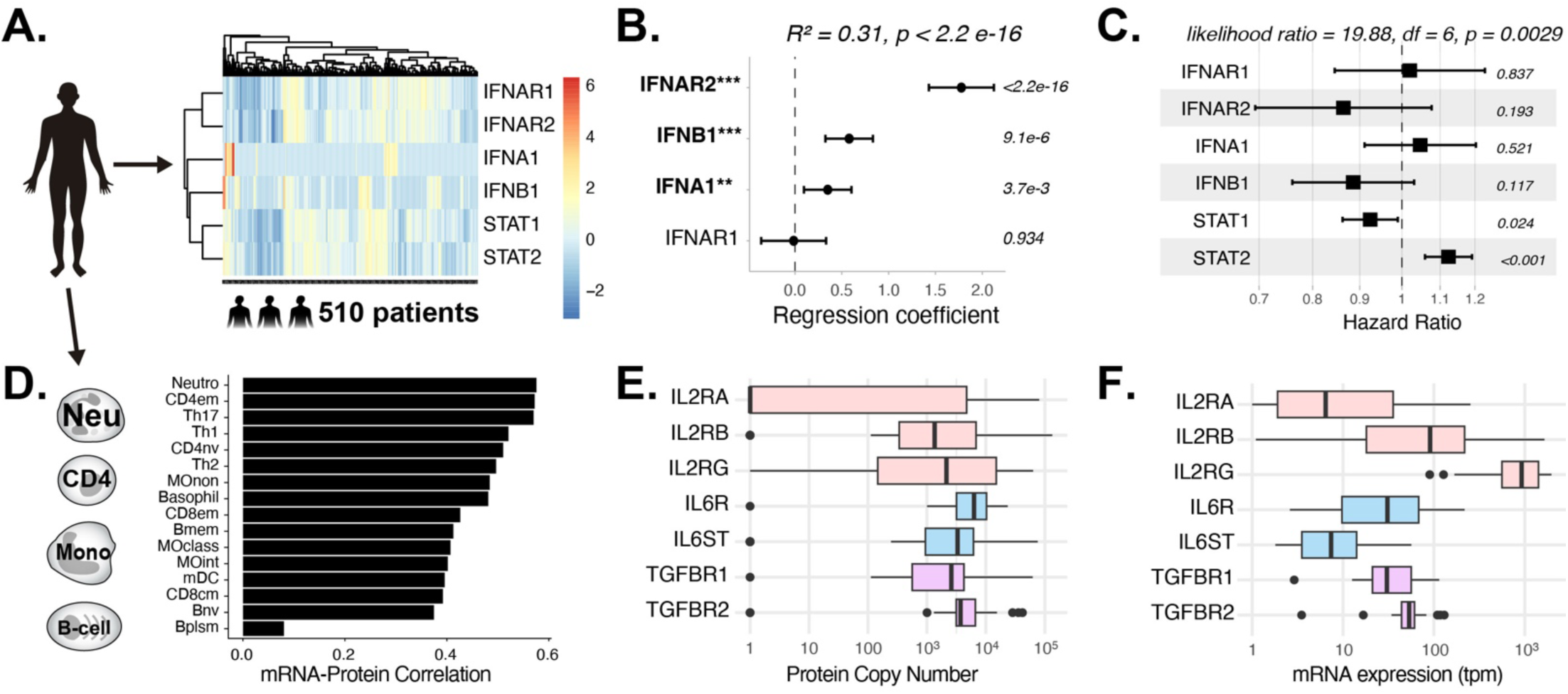
Clinical and protein-level validation of the cytokine avidity framework. <u>(A-C) Analysis of type I interferon signaling in</u> the TCGA lung adenocarcinoma cohort (n = 510).^74, 75^ **(A)** Expression of IFNA1, IFNB1, IFNAR1, and IFNAR2 and inferred STAT1 and STAT2 regulon activities across patients. **(B)** Multivariable linear regression of STAT1 regulon activity against ligand and receptor expression. IFNA1, IFNB1, and IFNAR2 showed significant independent associations with downstream signaling, and the combined model explained 31% of STAT1 activity variance (R^2^ = 0.31), consistent with the multivariable relationship predicted by Equation 7a. Points and error bars indicate regression coefficients and 95% confidence intervals; values indicate coefficient p-values. **(C)** Multivariable Cox proportional-hazards analysis of the complete type I interferon module, which was significantly associated with overall survival (p = 0.0029). <u>(D-F) Comparison of receptor mRNA and protein abundance across purified human immune-cell</u> populations.^60, 76^ **(D)** Pearson correlations between mRNA and protein abundance for 84 cytokine receptors across matched cell types. **(E-F)** Protein copy number (E) and mRNA abundance (F) for IL-2, IL-6, and TGFβ receptor systems. Relative receptor abundance was broadly conserved between modalities, including IL2RA, IL6ST, and TGFBR1 as the generally lower abundance receptor subunits.

## RESULTS

### Derivation and Validation of Avidity (EC_50_) Equations

#### Previous Solution Foundational Work

In 2008, Whitesides and colleagues derived an exact analytical solution for ternary complex equilibria in which a bivalent ligand (e.g. antibody) binds two copies of the same receptor.^52^ Building on this approach, in 2014 we derived an exact analytical solution for ligands that engage two distinct receptors (R_1_ and R_2_), as occurs in cytokine signaling.^51^ Together, these solutions provide analytical starting points for homo- and heterotypic ternary binding, alongside other theoretical treatments of multivalent binding at cell surfaces and interfaces.^39, 40, 55^ In the present work, we extend these frameworks by explicitly accounting for receptor confinement at the cell surface and deriving the corresponding EC_50_ relationships.

As detailed in the supporting information, we extend both frameworks by accounting for the reduction in translational entropy that occurs when receptors are confined to a cell surface,^56^ defining a restricted volume (V_surf_) in which surface-bound species interact.^56, 57^ A key result of this derivation is that EC_50_ depends entirely on the effective surface interaction volume (V_surf_) and is independent of bulk-solution volume (V_soln_), under the assumptions of this model. Accordingly, all concentration terms in the equations below carry a “surf” superscript. Throughout, [R]_t_ denotes total receptor concentration, K_1_ and K_2_ are equilibrium dissociation constants for R1 and R2, analogous to monovalent K_d_ values for each receptor interaction. In the monovalent limit (single receptor), this framework reduces to the classical relationship EC_50_ = K_d_. Although these equations describe receptor-complex formation, the predicted dependence on ligand affinity and receptor abundance is also observed in downstream signaling measurements. We distinguish below between binding EC_50_ and functional signaling EC_50_ and extend the equilibrium framework to steady-state conditions in which kinetic processes can shift potency.

#### Antibody Avidity Model

As detailed in the Supporting Information, we extended Whitesides’ homo-dimeric ternary complex model^52^ with our EC_50_ derivation and volume adjustment to obtain a closed-form expression for antibody binding at cell surfaces as a function of antigen density ([R]_t_^surf^) and monovalent binding constant (K_d_) (Eq. 4).

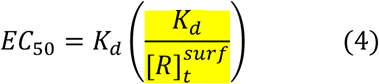

This equation reveals that avidity enhancement is governed by a single dimensionless ratio, K_d_/[R]_t_^surf^, providing a clear mechanistic explanation for how receptor density controls cellular sensitivity to multivalent ligands. Notably, the 10- to 1,000-fold potency enhancement of IgG over monovalent Fab fragments,^34–36, 46, 47, 49, 54^ widely cited in immunology textbooks,^58, 59^ emerges directly from this equation when evaluated across the typical range of copy numbers of cell-surface proteins (Fig 2 C-D).^60^ For the antibody formulation above, we assume independent equivalent binding sites (α = 1) as described in a cooperativity note section of supporting information (section 2.2).

An illustrative application is the long-standing challenge of broadly neutralizing antibodies against HIV. Studies have consistently shown that the avidity enhancement (IgG/Fab EC_50_ ratio) for HIV-targeting antibodies is only 1- to 10-fold, compared to 100- to 1,000-fold for antibodies against similarly sized enveloped viruses (Fig 2 C-D).^49, 54^ This disparity has been qualitatively attributed to the unusually low spike density on HIV virions, estimated at 10- to 100-fold fewer than comparable viruses based on cryo-EM studies (Figure 2C).^53^ Our framework explains this observation quantitatively: low [R]_t_^surf^ increases the K_d_/[R]_t_^surf^ ratio, compressing avidity enhancement into a narrow range. Furthermore, the equation predicts that increasing monovalent binding constant beyond the ~10 nM ceiling of natural affinity maturation^61^ can recover neutralizing potency, consistent with reports that exceptional target binding (sub-nanomolar K_d_) can restore broad neutralization against HIV.^62^

#### Cytokine Avidity Framework

In a similar manner, we extended our previous exact analytical solution to cell surfaces for ligands that bind two distinct receptors (R_1_, R_2_) with different affinities (K_1_, K_2_).^12, 51^ While we derive equations for the N-valent same-receptor case (antibodies) and both 2- and 3-valent heteromeric cases (cytokines), we focus here on the 2-receptor system as it is the most common and best supported by experimental data. Assuming R_1_ is the limiting receptor from a mass balance standpoint, we derive Equation 4, which includes a cooperativity factor (α) that accounts for stabilizing (α > 1) or destabilizing (α < 1) interactions between receptor subunits: ^51^

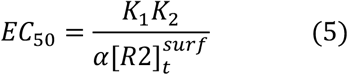

Conceptually, when one receptor is limiting from a mass-balance standpoint, its abundance constrained the maximum number of complete signaling complexes that can form, whereas an excess receptor can determine the efficiency of multivalent complex formation and therefore modulate EC_50_. Thus, receptor subunits can contribute asymmetrically to efficacy and potency. Which receptor occupies which role depends on the receptor stoichiometry of the cell-type or experimental context.

To validate our EC_50_ equations, we integrated data from biophysical studies, in vitro assays, murine single-cell RNA sequencing, and human clinical datasets (Fig 2-5). These relationships provide a common framework for interpreting how differences in receptor abundance can contribute to context-dependent differences in measured potency.

#### Quantitative Validation with Type-1 Interferons

We focus on the type I interferons throughout the remainder of this paper for three reasons. First, their receptors are expressed on nearly all nucleated cell types, providing a broad platform to assess signaling across diverse biological contexts.^58, 64^ Second, the type I interferon family includes approximately 20 natural and engineered ligands that all signal through the same receptor pair (IFNAR1, IFNAR2), differing only in their K_1_ and K_2_ values, making them an ideal test case for exploring how changes in binding constant and receptor expression shape potency.^63^ Third, type I interferons are among the most extensively studied cytokines biophysically, with comprehensive K_d_ measurements available across all variants, enabling rigorous quantitative application of our framework in a way not yet possible for other cytokine families.^20, 21^

We began our validation with *in vitro* cell-based assays, which provide a uniquely controlled setting to quantify both *<u>binding EC_5_</u>_0_* and *<u>functional signaling EC_50_</u>* through dose-response analysis (Fig 3C). The most comprehensive biophysical data come from the Schreiber and Piehler groups, who systematically measured type I interferon binding affinities, cell-surface EC_50_’s, and downstream responses including STAT1 phosphorylation and antiproliferative activity. ^21–23, 29, 63, 65, 66^

A key finding from this body of work was that cellular EC_50_’s correlated more strongly with the product of the two interferon binding affinities (K_1_ x K_2_) than with either K_d_ alone: an empirical observation that our model explains mechanistically.^20, 21^ Figure 3C compiles 42 binding affinities across 21 natural and engineered type I interferons, paired with EC_50_’s for both direct cell-surface association and downstream antiproliferative signaling. The relationship was observed for both *<u>cell-surface binding</u>* and *<u>downstream antiproliferative responses</u>*, indicating that the affinity dependence predicted for complex formation remains detectable at the level of functional signaling. ^20, 21, 63^ Furthermore, Figure 3D highlights the quantitative relationship between EC_50_ and the measured surface density of IFNAR1, which was present in excess relative to its signaling partner under the conditions of these experiments, confirming the additional importance of receptor expression.^23^

#### Extension from Equilibrium Binding to Steady-State Signaling

The EC_50_ equations derived above assume pre-equilibrium conditions, in which assembly and disassembly of the ligand-receptor complex occur more rapidly than downstream biological processes.^57, 67^ Under this approximation, binding potency can be described directly from equilibrium binding constants and receptor abundance. However, as illustrated in Figure 3C, the *EC_50_ for receptor binding* is rarely equal to the *EC_50_ of a downstream signaling response*.^57^ When the lifetime of the signaling complex approaches the timescale of processes such as signal generation or ligand-induced receptor endocytosis, these downstream rates can impose an additional kinetic constraint on apparent potency.^61, 68^

We therefore extended the equilibrium derivations to steady-state models in which formation and loss of the functional signaling complex are balanced over the timescale of the biological response (Supporting Information section 3). As in classical Michaelis-Menten kinetics, the resulting signaling EC_50_ contains an additional kinetic term determined by the relative rates of complex formation and downstream processing (Figure S11).^69^ This creates a kinetic limit beyond which further improvements in equilibrium binding affinity produce progressively smaller changes in functional potency.^61^ Importantly, the corresponding multivalent steady-state solutions retain the general structure of the equilibrium models, with ligand concentration, receptor abundance, and multivalent receptor engagement continuing to determine signaling sensitivity, while the downstream kinetic term shifts the effective EC50 (Table S1), exactly as is observed in Figure 3C.

### Extension to In Vivo and Transcriptomic Cytokine Signaling

The preceding analyses test the avidity relationships in controlled systems where binding affinities and ligand concentrations are experimentally defined. Applying these relationships *in vivo* requires a different formulation because ligand abundance, receptor expression, and downstream signaling vary simultaneously across cells and tissues. We therefore embedded the derived EC_50_ relationship (eq 5) within a Hill-type response model (eq 1) ^42, 43^ and rearranged it into forms suitable for statistical analysis of transcriptomic data, where SIG represents down-stream signaling quantified by transcription factor target gene-<u>sig</u>natures:

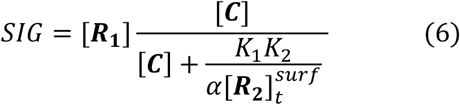

Under sub-saturating conditions ([C] << EC_50_) and taking the logarithm of both sides yields a linear signaling model in which all variables contribute:

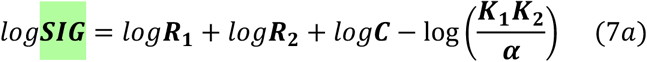

Conversely, under saturating conditions (C >> EC_50_), potency-dependent terms become negligible and signaling is determined primarily by the limiting receptor:

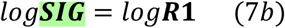

Conceptually, this separation arises because the limiting receptor (R1) constrains the total number of signaling complexes that can form, limiting total signaling flux. Conversely, the excess receptor governs how efficiently the ligand is captured into ternary complexes, thereby modulating EC_50_. This division of roles is not apparent in classical monovalent models but emerges naturally from mass-balance constraints in multi-valent systems.

This log-scale formulation of these equations is naturally suited to transcriptomic data, where gene and protein expression are approximately log-normally distributed.^70^ Critically, these forms are immediately amenable to statistical inference, enabling multivariable regression to identify which sources of ligand or receptor variance most strongly drive signaling across cell types and disease contexts.

#### Experimental validation

To validate our framework in vivo, we leveraged a dataset from Cui et al. in which single high doses of 86 cytokines were administered to mice and transcriptional responses were measured by scRNAseq across 18 immune cell types from draining lymph nodes (Fig 4A).^71^ Because cytokines were administered at high dose, this experiment approximates the saturating regime of Equation 7b, in which variation in ligand potency becomes less influential and signaling is predicted to depend primarily on the abundance of the limiting receptor. Consistent with this prediction, baseline Ifnar1 expression strongly correlated with the magnitude of the type I interferon transcriptional response magnitude across immune cell types, while Ifnar2 did not (Fig 4B). Across the 16 cytokines, receptor subunits frequently showed asymmetric associations with response, with one receptor typically displaying a stronger positive correlation than its partner (Fig 4C-D), a pattern consistent with the predicted receptor asymmetry.

We next asked whether the broader ligand-receptor relationships predicted under non-saturating conditions could be detected in human tissues, where ligand abundance, receptor expression, and downstream signaling vary simultaneously.

#### Spatial validation in human lung cancer

To determine whether the avidity framework could be applied to human transcriptomic data, we first analyzed the original CosMx Spatial Molecular Imaging dataset from a human lung adenocarcinoma specimen (Fig 1B, S1).^10^ This dataset contains single-cell spatial transcriptomic measurements across approximately 100,000 cells. Because cytokines can act on neighboring cells, cytokine expression was estimated from the surrounding 50-µm spatial niche, whereas receptor expression and downstream signaling were treated as cell-intrinsic properties. Downstream signaling was quantified using two complementary approaches: Hallmark transcriptional signatures from MSigDB^72, 73^ and transcription-factor target regulons compiled by DoRothEA.^70^ We then tested whether ligand and individual receptor-subunit expression predicted downstream signaling as expected from Equation 7a. The two signaling readouts produced broadly concordant results, but the relative contributions of individual signaling components differed substantially among cytokines (Fig S2-S3). Multivariable analysis identified four systems (IFNα, IL-2, TNF, and TGFβ) in which multiple ligand or receptor components contributed independently to downstream signaling (Fig S4). Combined models explained 48%, 73%, 29%, and 68% of signaling variance, respectively, consistently exceeding the predictive value of individual components. Thus, cytokine signaling within intact human tissue was better explained by joint ligand-receptor relationships than by any single ligand or receptor variable alone.

#### Population Level Validation and Clinical relevance

We next asked whether these relationships generalized beyond a single spatially profiled tumor by analyzing bulk RNAseq from 510 patients in the TCGA Lung Adenocarcinoma PanCancer Atlas cohort.^74, 75^ We evaluated the four cytokine systems identified above using their canonical downstream transcription-factor regulons: STAT1 for type I interferon, STAT5A for IL-2, SMAD3 for TGFβ, and NFKB1 for TNF. Despite the substantial differences between spatial and bulk transcriptomic measurements, multiple ligand and receptor components independently predicted downstream regulon activity for all four cytokine systems validated in the CosMx dataset (Figure 5A-B, Figure S5). For type I interferon, IFNA1, IFNB1, and IFNAR2 each contributed independently to STAT1 activity, and the combined model explained 31% of variance across patients (R^2^ = 0.31; Fig 5B). We separately assessed clinical relevance using multivariable Cox proportional-hazards models incorporating the complete ligand-receptor-response axis. All four cytokine modules were associated with overall survival by likelihood-ratio testing (Fig 5C, S5); for type I interferon, the six-variable model was significant (likelihood-ratio = 19.88, df = 6, P = 0.0029). These survival analyses provide an orthogonal measure of clinical relevance rather than a direct validation of the mechanistic avidity equations.

#### Independent validation of mRNA-to-protein correspondence

Finally, application of the framework to transcriptomic data assumes that relative receptor mRNA abundance provides useful information about receptor protein abundance (Fig 1A). We evaluated this assumption independently using transcriptomic and quantitative proteomic measurements from purified primary human immune-cell populations from healthy donors.^60, 76^ Across 84 cytokine receptors represented in matched cell populations, receptor mRNA and protein abundance showed an overall positive correspondence of approximately r = 0.5 (Fig 5D). Relative differences among receptor subunits were also broadly conserved for representative cytokine systems, including IL2RA, IL6ST, and TGFBR1 as generally lower-abundance receptors. Quantitative proteomics further showed that many cytokine receptors are present at thousands to tens of thousands of copies per cell, while some receptor/cell-type combinations occur at substantially lower abundance (Fig S6).

Together, these analyses test three complementary aspects of transcriptomic application of the framework. Spatial transcriptomics demonstrates that ligand and receptor abundance jointly predict downstream signaling within intact human tissue; TCGA shows that these relationships are conserved across patients and associated with clinical outcome; and independent primary immune-cell measurements support mRNA abundance as an informative, although imperfect, surrogate for receptor protein abundance. Broader validation across additional diseases and clinical settings will remain important.

## DISCUSSION

### Analytical Models and Receptor Asymmetry

This study develops an avidity-based framework for understanding how multivalent receptor engagement alters ligand potency at cell surfaces. Unlike monovalent interactions, in which potency is primarily determined by a binding constant, the EC_50_ of a multivalent interaction also depends on receptor abundance. For multicomponent cytokine receptors, individual subunits can play asymmetric roles: one can constrain the maximum number of signaling complexes that form, whereas another controls the efficiency of multivalent complex formation and therefore cellular sensitivity. These relationships explain established observations including density-dependent enhancement of antibody binding and receptor-dependent changes in type I interferon potency. ^21–23, 29, 63, 65, 66^

This asymmetry is particularly relevant because receptor expression varies substantially across cell types and cellular states. Cytokine sensitivity can therefore change without altering the cytokine or its intrinsic binding affinities. Which receptor most strongly affects sensitivity depends on receptor stoichiometry and ligand concentration, providing a quantitative basis for longstanding observations that variation in a single receptor subunit can dominate responsiveness even when multiple receptor components are required.^12–16, 25, 27–31^ Receptor context thus provides one mechanism through which the same extracellular cytokine can generate different responses across cell types.

The *equilibrium binding EC_50_* derived here should not, however, be assumed to equal the *EC_50_ of a downstream biological response* (as illustrated in Figure 3C). Signaling, receptor endocytosis, degradation, recycling, and related processes can alter functional potency when they occur on timescales comparable to receptor-complex assembly.^61, 67^ Our steady-state extensions account for these effects through an additional kinetic constraint while retaining the underlying dependence on ligand concentration, receptor abundance, and multivalent engagement. The equilibrium framework should therefore be viewed as a mechanistic baseline that can be incorporated into more detailed ODE, stochastic, or signaling-network models when greater temporal resolution is required.

### Transcriptomic methods

A second contribution of this study is translating these molecular relationships into a form that can be evaluated using transcriptomic data. Ligand and individual receptor-subunit abundance jointly predicted downstream transcriptional activity in human spatial transcriptomic data, and related multivariable relationships were observed across 510 lung adenocarcinoma patients. The corresponding ligand– receptor–response modules were also associated with overall survival, supporting their clinical relevance without implying that the avidity mechanism itself determines outcome. Quantitative proteomics from primary human immune cells further showed that receptor mRNA abundance is an informative, although imperfect, surrogate for relative protein abundance. Together, these analyses suggest that the molecular dependencies predicted by the model remain detectable across increasingly heterogeneous biological systems.

The framework is complementary to existing cell-cell communication methods such as CellChat and CellPhoneDB as illustrated in Figure S7. ^4–6^ These methods primarily identify and score candidate communication between sender and receiver populations, whereas our approach asks whether ligand abundance, individual receptor subunits, and downstream responses follow relationships predicted from multivalent binding. This distinction is especially important for multi-subunit receptors: CellChat, for example, combines receptor components using a geometric mean, whereas the avidity framework allows individual receptor subunits to contribute differently to potency and maximal response. Thus, communication methods identify where signaling may occur, while the present framework provides a mechanistic hypothesis for how receptor context modifies signaling sensitivity.

### Limitations & Future work

Several assumptions define the framework’s range of applicability. First, it assumes a well-mixed cell surface governed by conventional mass-action relationships; strong receptor clustering,^55^ membrane compartmentalization, immobilized ligands, or organized interfaces such as immunological synapses may require explicitly spatial models.^77–79^ Second, the deterministic formulation assumes sufficiently large molecule numbers, whereas low-copy receptor systems may require stochastic treatment^80^ and very high surface densities may introduce non-ideal effects.^81^ Third, transcriptomic applications are limited by imperfect correspondence between mRNA and surface protein abundance^60, 70^ and by incomplete knowledge of binding constants and cooperativity parameters. When independent biophysical measurements are unavailable, the cooperativity factor α is therefore best interpreted as part of an effective affinity term rather than as a separately identifiable parameter.^18, 52, 82^ Approaches that propagate uncertainty in expression, affinity, and kinetic parameters may be useful as these models are expanded.^83, 84^

The analytical framework is also readily extensible. Mechanistically informed EC_50_ relationships could be incorporated into platforms such as CellChat (Fig S7),^4, 85^ VCell,^79^ SimBiology,^86^ SMART,^78^ or NetFlux,^87^ particularly where Hill parameters are currently selected empirically. Future extensions could explicitly include spatial diffusion,^7, 8^ receptor trafficking,^3, 88^ stochastic low-copy interactions,^80^ and higher-valency antibody systems such as IgA and IgM.^89^ The value of the present formulation is not that it captures every kinetic or spatial feature of signaling, but that it provides a tractable mechanistic baseline connecting binding affinity, receptor abundance, and multivalent complex formation.^90^ In this way, avidity provides a quantitative bridge between molecular binding and cell-type-specific signaling across complex biological systems.

## Computational Method Details

### Analytical Model Derivations

All analytical models were derived using the laws of mass action and conservation of mass and either equilibrium conditions (S01-S69) or steady-state conditions (S70-S134). For the antibody and cytokine avidity models we adjusted the receptor conventions to match the soluble-system models from which they were derived. Specifically, Whitesides^52^ and our early work^51^ treat “B” as the bridging species (antibody or cytokine), with receptors labeled as A-B-A for the homodimeric antibody case (A = antigen) and A-B-C for the heterodimeric cytokine case (A = R1 and C = R2). Figures S8-S9 illustrate the key difference between previously published exact analytical models and the models presented here in: this current work explicitly treats the immobilization volume at the cell-surface following antibody/cytokine binding (Figure S10, equations S01-02). EC_50_ equations were obtained by solving for the antibody or cytokine concentration at which 50% of maximum complex is formed; additional proofs are detailed in our 2014 JACS paper.^51^ This approach was applied to heterodimerizing ligand receptors (A-B-C) such as the type-1 interferons in equations S3-S18, homo-dimerizing receptors (A-B-A) for bivalent IgG in equations S19-S35, homo-trimerizing receptors (A_3_B) for cytokines like TNF in equations S36-S52 and hetero-trimerizing receptors (A-B-C-D), for cytokines like Interleukin-2 in equations S53-S69.

### Literature Dataset and Database Curation

All experimental data analyzed in the paper was obtained from manual curation of the primary literature (e.g. major cytokine definitions,^3, 4^ Figure 3C,^21, 63, 66^ Figure 3D,^22, 23^) and is publicly available in our GitHub Repository (https://github.com/fingolfn/avidity-math). In addition, all data was obtained from either the original publications: antibody avidity^49, 54^ cytokine affinities, binding and signalling,^21–23, 29, 63, 65, 66^ in vivo murine scRNAseq,^71^ and spatial transcriptomics.^10^ Finally, raw data along with visualization code is available through a GitHub Repository (https://github.com/fingolfn/avidity-math).

### Data Analysis and Visualization

All data processing, statistical analysis, and visualization were performed in R statistical software (version 4.5.1). Heatmaps were generated using the gplots and RColorBrewer packages, and pathway analyses were performed using VIPER and msigdbr. All analysis scripts and visualization code are publicly available through a GitHub repository (https://github.com/fingolfn/avidity-math).

### Statistical Analysis

All statistical analyses were performed in R (version 4.5.1). Correlation analyses used Pearson correlation coefficients with two-tailed significance testing unless otherwise specified. Linear regression analyses were performed using the base R lm function on log10-transformed variables where appropriate. Coefficients of determination (R^2^), correlation coefficients (r), and associated P values were calculated directly from the fitted models. Bootstrap confidence intervals were estimated using 1,000 resampling iterations. Heatmaps display scaled correlation coefficients or enrichment scores as indicated in the corresponding figure legends. Statistical significance was defined as P < 0.05. Exact statistical tests, sample sizes, and error bar definitions are described in the relevant figure legends and specific figure methods subsections below.

### Soluble-ligand avidity analysis (Figure 3)

Binding affinities (Kd) and functional EC_50_ values for type I interferon variants were digitized from published biochemical and cell-based assays.^21–23, 29, 63, 65, 66^ To allow direct comparison across ligands, all K_d_ and EC₅₀ values were scaled to the IFN-α2 reference. For each variant we calculated the product of its IFNAR1 and IFNAR2 affinities (K1 × K2). Log10-transformed EC_50_ values were regressed against log10(K1 × K2) with the base **lm** function in R; slope, intercept, r, and two-tailed *p* were annotated on the plot. A second data set linking IFNAR1 surface copy number to EC₅₀ was analyzed in the same manner, with replicate error propagated as orthogonal error bars drawn with arrows().

### Exogenous cytokine with murine scRNAseq^71^ (Figure 4)

single-cell RNAseq from draining lymphnodes^71^ were aggregated into TPM-based pseudobulk matrices (PBS_data_filtered). Gene-level cytokine responsiveness z-scores (Relative to PBS controls) were extracted from stimulation experiments (cell_responsiveness). A hand-curated list of 16 cytokines was mapped to their signaling receptors with a lookup table (cytokine_receptors_SIMPLE.csv). Pearson correlation coefficients were computed between every receptor-cytokine pair with cor() and displayed as heatmaps using the gplots or pheatmap packages. For key axes (e.g., IFNAR1/2 vs. IFNA1 or IFNB1 response) individual linear fits were over-plotted to quantify goodness-of-fit. Monocytes were excluded from the interferon correlation analysis due to absent variance in transcriptional response across the profiled interferons. Because these receptor-response correlations were evaluated across multiple cytokines without adjustment for multiple testing, they were treated as exploratory and interpreted primarily based on the pattern and magnitude of association rather than formal statistical significance.

### Ligand, multi-valent receptor, transcription factor networks (S2-S7)

We defined a curated network of 19 cytokine ligand-response axes with established multicomponent receptor engagement or receptor oligomerization^17, 19^ and canonical downstream transcription-factor responses.^3^ These included IFNG, IFNA1, IFNB1, IL2, IL4, IL6, IL7, IL10, IL12A, IL15, IL17A, IL18, IL21, IL22, IL23A, TNF, TGFB1, CSF2, and CSF3. Ligand–receptor assignments were derived from CellChat and curated literature,^4, 85^ while downstream transcription-factor relationships were defined from established signaling pathways.^3^ For each cytokine system, the corresponding receptor subunits were extracted from the CellChat interaction table and filtered to those present in the CosMx 1,000-gene panel. One annotation error was corrected: IL20RA was reassigned to IL2RA for the IL-2 signaling system. Six cytokine systems were retained for analysis based on the availability of all three components (ligand, receptors, and downstream signaling readout) in the CosMx panel:^10^ IFNα, IFNγ, IL-2, IL-6, TNF, TGFβ, and their associated receptor subunits.

### CosMx Samples: Neighborhood definition for spatial pseudobulk analysis (Fig S1-S4):^10^

Spatial x-y coordinates and cell-type annotations exported from CosMx SMI were merged to construct a “local composition” matrix for every cell. For each index cell we identified all neighboring cells whose centroids fell within a 50 µm Euclidean radius. The counts of each canonical cell type within that disc were converted to fractions of the total neighborhood and stored as a row vector, yielding an *N x C* matrix (cells x cell-type fractions). This matrix was logit-transformed to stabilize variance, scaled to unit variance per column, and subjected to *k*-means clustering with *k = 6* (nstart = 100) to obtain six reproducible neighborhood classes (k1-k6). We selected k = 6 to provide a coarse-grained representation of the major tumor, stromal, myeloid, and lymphoid microenvironments while maintaining sufficient observations within each cell-type × niche group for downstream pseudobulk analysis. For downstream analyses we summed raw gene-level counts for all cells sharing a common label (cell-type_neighborhood), generating pseudobulk expression profiles that capture both cellular identity and micro-environmental context. These profiles served as the input for the VIPER-algorithm based pathway enrichment described below.

To reflect that signaling is driven by the local microenvironmental cytokine concentration rather than a cell’s own ligand transcripts, cytokine expression was assigned at the niche level while receptor expression and downstream signaling were treated as cell-intrinsic. For each niche, a weighted average of gene expression was computed by multiplying the cell-type frequency composition of the niche (derived from the neighborhood clustering above) by the cell-type-level pseudobulk expression matrix. The resulting niche-level ligand estimates were then propagated to all cell-type x niche pseudobulk groups within that niche.

### CosMx Samples: Pathway-level inference in spatial neighborhoods (Fig S2A)

To put receptor– response relationships into pathway context, hallmark gene-set activities were inferred with the VIPER algorithm.^91^ Human hallmark sets were downloaded with msigdbr,^73^ stripped of the “HALLMARK_” prefix, and converted into regulons assuming unitary tf-mode and likelihood. VIPER was run on both cell-type-level pseudobulk (logRPM) and niche-level logTPM matrices with a minimum regulon size of 10 genes. The resulting enrichment scores (for example “IL2_STAT5_SIGNALING” and “IL6_JAK_STAT3_SIGNALING”) were regressed against log-scaled receptor abundance for all niches.

### CosMx Samples: Transcription factor target signatures (Fig S2B)

As a complementary signaling readout to Hallmark gene signatures, we computed transcription factor (TF) target activity scores using Dorothea-compiled regulons^92^ via VIPER.^91^ Each cytokine was mapped to its canonical downstream transcription factor(s) based on established signaling cascades: IFNα/β → STAT1, STAT2, IRF9; IFNγ → STAT1; IL-2/IL-7/IL-15 → STAT5A/B; IL-6/IL-10/IL-21/IL-22/IL-23 → STAT3; IL-4 → STAT6; IL-12 → STAT4; TNF/IL-17/IL-18 → NFKB1, RELA, JUN, FOS; TGFβ → SMAD2, SMAD3. TF-target activity was scored as the VIPER enrichment of each TF’s regulon, providing a mechanism-based measure of downstream signaling independent of the phenotypically defined Hallmark signatures.

### CosMx Validation of Multi-variate Cytokine Signaling (Figure S2-S4)

For each of the eight cytokine systems, univariate linear regressions were performed between downstream signaling (Hallmark or TF-target activity score) and each individual predictor variable (niche-level ligand expression or cell-intrinsic receptor expression). R^2^ values and 95% confidence intervals were estimated by bootstrap resampling (R = 1,000 iterations). For visualization and comparison across signaling frameworks (Figure 5), predictor variables were ranked by the average R^2^ across both Hallmark and TF-target readouts (Fig S2). Multivariable linear models were then fit for each cytokine system using all available ligand and receptor variables as predictors. Independent contributions of individual predictors were assessed via coefficient significance in the multivariable fits. Cytokine systems for which multiple predictors showed significant independent contributions (IFNα, IL-2, TGFβ, TNF) are reported in the supporting information (Fig S4). To ensure robust pseudobulk estimates, cell-type × niche combinations were included in regression analyses only if the cell type constituted >5% of the total cells within that niche. This threshold excluded rare populations for which pseudobulk expression estimates would be unreliable due to low cell counts.

### TCGA Clinical Validation of Multivariate Cytokine Signaling (Figure 5, S5)

To independently evaluate the multivariable signaling relationship predicted by Equation 7a and assess its clinical relevance, we analyzed bulk RNA-seq and clinical data from the TCGA lung adenocarcinoma Pancancer Atlas cohort (n = 510 patients).^75^ RSEM-normalized gene expression values were log10-transformed and standardized across patients for each gene. Transcription factor (TF) activity was inferred from bulk RNAseq using VIPER and DoRothEA regulons restricted to confidence levels A-C (i.e. direct evidence of target gene regulation).

Four cytokine systems previously showing multivariable regulation in the CosMx analysis were evaluated: type I interferon, IL-2, TGFβ, and TNF. Multiple linear regression was used to test whether downstream TF-regulon activity could be jointly explained by expression of the corresponding ligand and receptor subunits, as predicted by Equation 7a (Fig S5). Models were fit for STAT1 activity using IFNA1, IFNB1, IFNAR1, and IFNAR2; STAT5A using IL2, IL2RA, IL2RB, and IL2RG; SMAD3 using TGFB1, TGFBR1, and TGFBR2; and NFKB1 using TNF, TNFRSF1A, and TNFRSF1B. Regression coefficients and 95% confidence intervals were used to assess the independent contribution of each signaling component.

Clinical relevance was assessed independently using multivariable Cox proportional-hazards regression with overall survival as the endpoint. For each cytokine system, models incorporated ligand expression, receptor-subunit expression, and downstream TF-regulon activity. Hazard ratios with 95% confidence intervals and model significance were calculated using the survival package in R. These survival analyses were used to determine whether variation across the complete ligand-receptor-signaling module was associated with patient outcome, rather than as a direct validation of the mechanistic avidity model.

### Cross-modal validation of mRNA as a surrogate for receptor protein abundance (Figure 5, S6)

To assess the use of transcript abundance as a surrogate for receptor protein abundance, we compared independently generated RNAseq and quantitative proteomic profiles of purified primary human immune-cell populations. RNAseq data were obtained from Monaco et al,^76^ who profiled 29 flow-sorted peripheral immune-cell populations from healthy donors, with expression represented as transcripts per million (TPM). Quantitative proteomic data were obtained from Rieckmann et al,^60^ who profiled flow-sorted human hematopoietic populations by high-resolution mass spectrometry and estimated protein copy numbers using the proteomic-ruler approach. Cell-type annotations were harmonized between datasets, proteomic replicates were averaged within each cell type, and analyses were restricted to genes and cell populations represented in both datasets. Because intracellular protein measurements may not accurately represent secreted ligand concentrations, comparisons were restricted to receptor genes contained within the CellChat cytokine networks.^4, 85^ RNA and protein abundance were transformed as log10(x + 1). For each receptor, Pearson correlation was calculated between mRNA and protein abundance across matched immune-cell populations. In addition, mean mRNA and protein abundance across cell types were calculated for each receptor and compared using Pearson correlation and simple linear regression. Representative cytokine receptors were additionally visualized by comparing their mRNA and protein abundance distributions across immune-cell populations.

## Lead Contact

Further information and requests for resources should be directed to and will be fulfilled by the Lead Contact, Eugene Douglass.

## Materials Availability

This study did not generate new unique reagents.

## Data Availability

The datasets analyzed during the current study are available from the original publications cited herein and from the GitHub repository: https://github.com/fingolfn/avidity-math.

## Code Availability

All custom R scripts used for data processing, statistical analysis, figure generation, and modeling are publicly available at: https://github.com/fingolfn/avidity-math.

## Author Contributions

E.F.D. conceived the study, developed the analytical framework, performed analyses, and wrote the manuscript. W.B. contributed computational analysis and manuscript editing. J.P.M. contributed conceptual guidance and manuscript editing. All authors approved the final manuscript.

