## Supplementary Information for "A Comprehensive Mathematical Model of Avidity in Cytokine Signaling"

**1.4 Clinical Validation of Avidity Models Assumptions leveraging Quantitative Proteomics of PBMCs** 7

### 1. Supporting Figures

#### 1.1 Background on single-cell spatial transcriptomic technology: CosMx

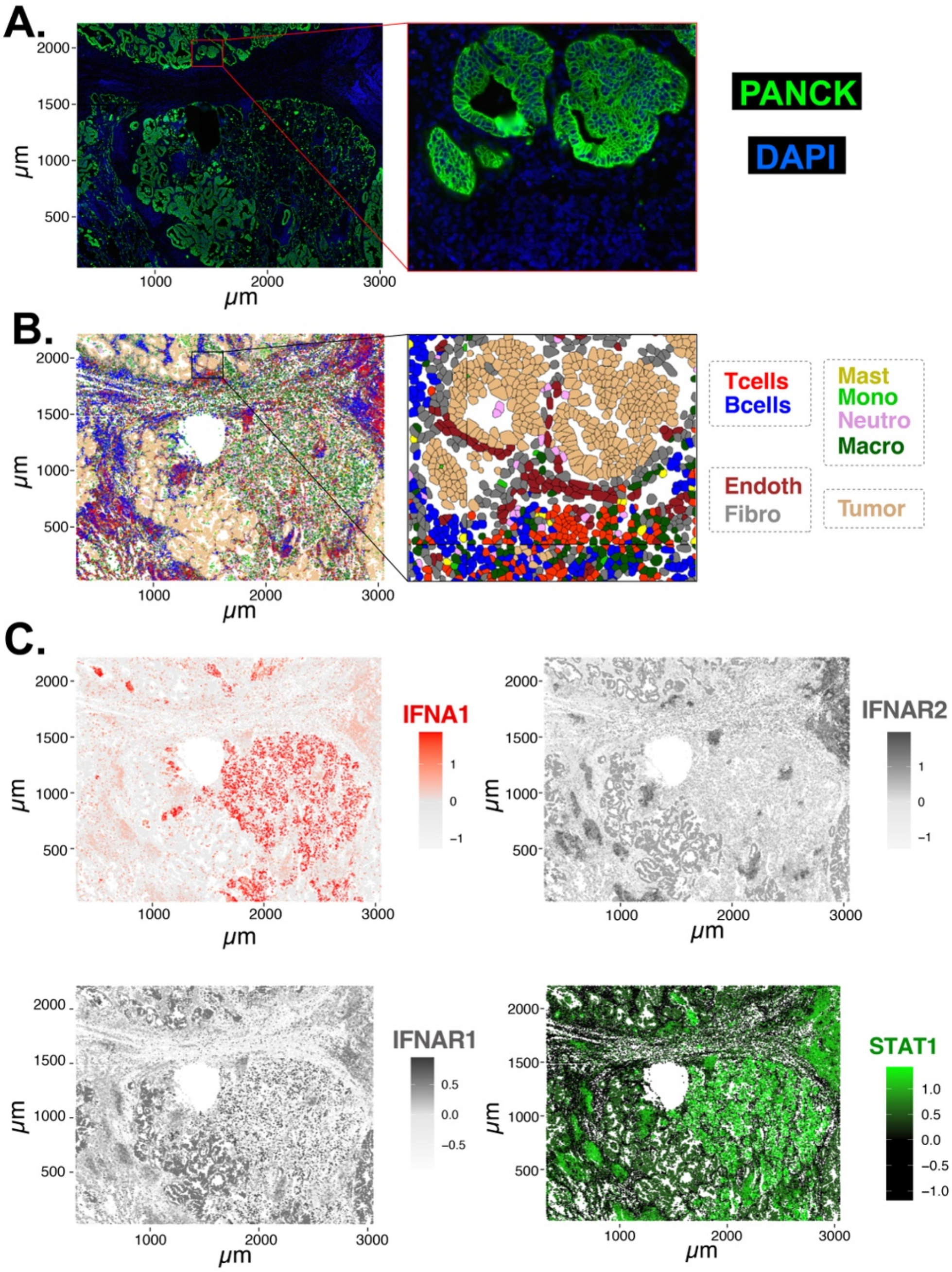

**Figure S1. Overview of the first published CosMx Spatial Molecular Imaging (SMI) dataset. This dataset, derived from a lung adenocarcinoma FFPE sample is the most commonly used benchmark for single-cell resolution spatial transcriptomic methods for the CosMx technology (A)** Immunofluorescence of an FFPE non-small cell lung cancer biopsy (PanCK, green; DAPI, blue). **(B)** nine cell types identified from the first published CosMx transcriptomic profiling. **(C)** Spatial maps of type I interferon signaling across the tissue section, illustrating IFNA1 ligand expression (top left), IFNAR1 (bottom left) and IFNAR2 (top right) receptor expression, and STAT1 target gene activity (bottom right). The spatial heterogeneity of all three components across cell types and tissue regions motivates a quantitative framework that integrates ligand, receptor, and signaling evidence.

#### 1.2 Clinical Validation of Avidity Model Across 100,000 single-cells (CosMx)

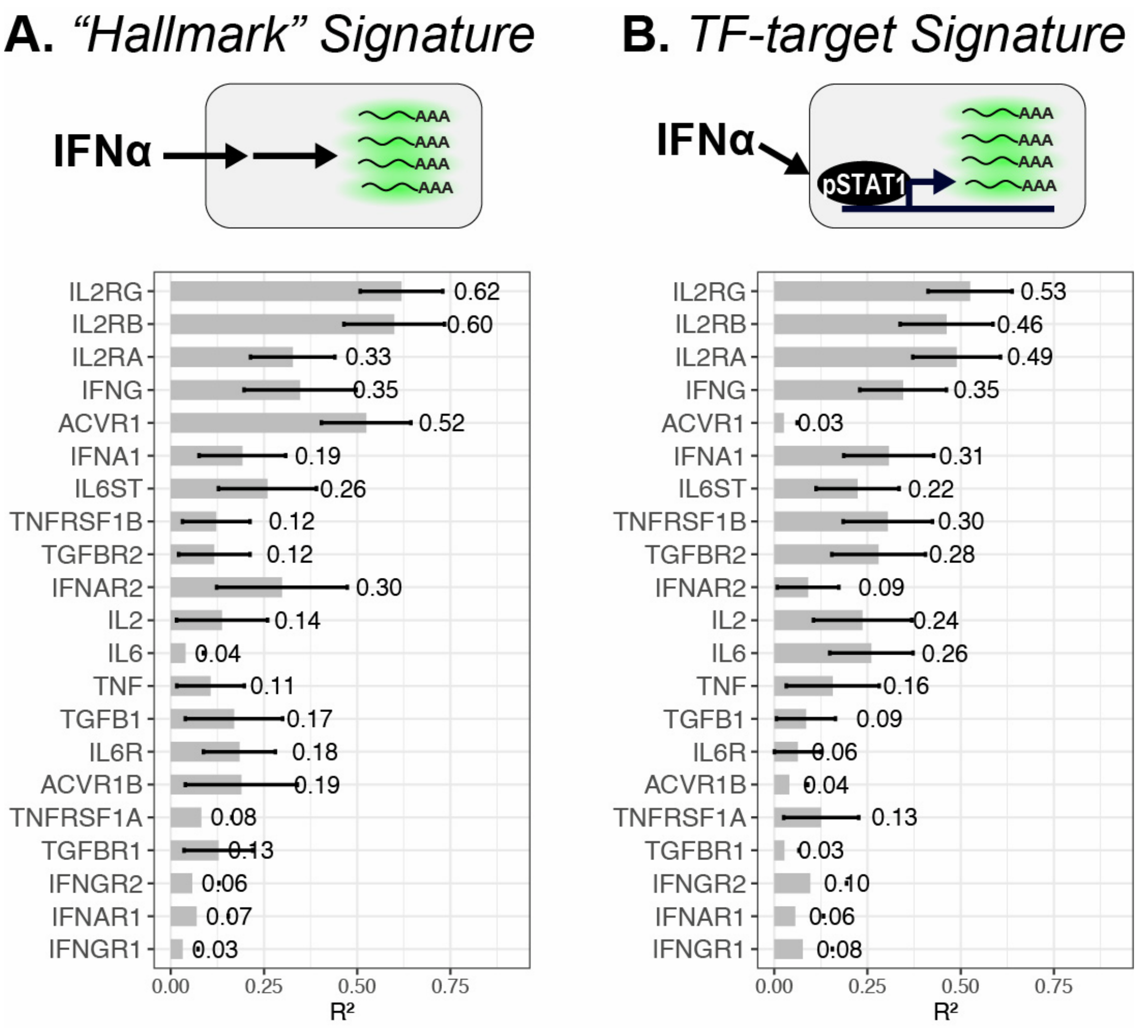

**Figure S2. Univariate prediction of downstream cytokine signaling from individual ligand and receptor variables in human NSCLC spatial transcriptomic data.**[1] Each bar represents the $R^2$ from univariate regression of downstream signaling against a single ligand or receptor variable across all cells in the CosMx dataset, aggregated across eight cytokine systems for which ligands, receptors, and signaling readouts are all available in the panel. Results are shown for two complementary signaling readouts: (A) Hallmark gene signatures from MSigDB and (B) Dorothea-compiled transcription factor target regulons (illustrated here for IFNα → pSTAT1). The two approaches are generally concordant, identifying receptor subunits such as IL2RG, IL2RB, and ACVR1 as strong predictors of signaling while other variables contribute minimally. *Statistical analysis*: Univariate linear regressions were fit across cell-type × niche pseudobulk groups using the lm function in R. $R^2$ indicates the proportion of downstream signaling variance explained by each individual predictor; 95% confidence intervals were estimated by bootstrap resampling (1,000 iterations). P values were not corrected for multiple comparisons, and these analyses should therefore be interpreted as exploratory

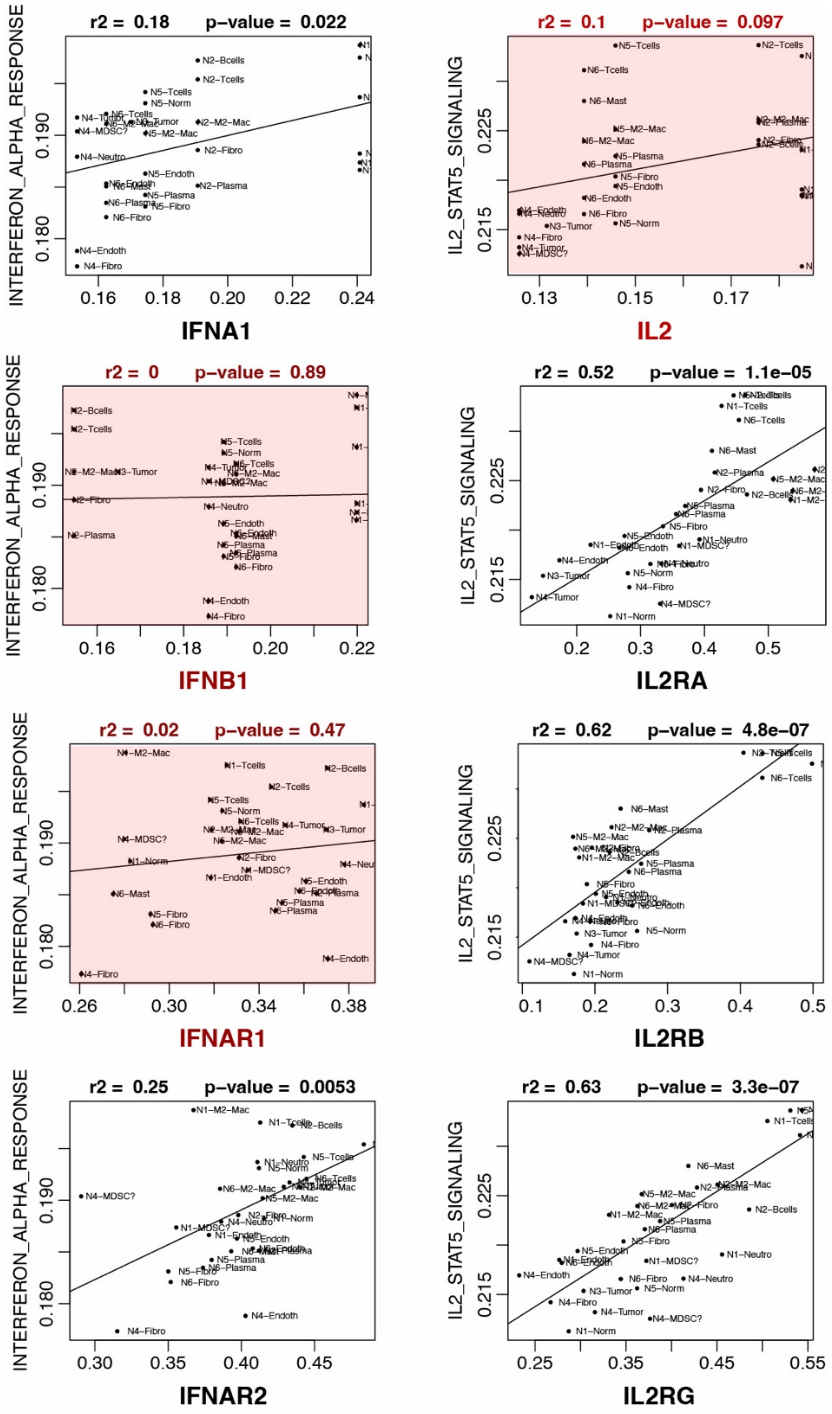

**Figure S3. Representative univariate regressions underlying the summary analysis in Figure 5**. Each panel shows the relationship between a single predictor variable (x-axis) and downstream signaling activity (y-axis) across pseudobulk groups defined by niche x cell-type (e.g., N5-Tcells), capturing both cell-intrinsic and spatial contributions to signaling. Left column: IFNα signaling (Hallmark INTERFERON_ALPHA_RESPONSE) regressed against niche-level IFNA1 ligand expression (R2 = 0.18), IFNB1 (R2 = 0.00), IFNAR1 (R2 = 0.02), and IFNAR2 (R2 = 0.25). Right column: IL-2 signaling (Hallmark IL2_STAT5_SIGNALING) regressed against niche-level IL2 ligand expression (R2 = 0.10) and cell-intrinsic receptor expression for IL2RA (R2= 0.52), IL2RB (R2 = 0.62), and IL2RG (R2 = 0.63). Red shading and text indicate non-significant associations (p > 0.05). For IFNα, ligand and IFNAR2 show stronger associations than IFNAR1; for IL-2, all three receptor subunits showed nominally significant associations while niche-level ligand is not. These patterns are consistent with the aggregated R2 values reported in Figure 5. *Statistical analysis:* Each relationship was evaluated by univariate linear regression across cell-type × niche pseudobulk groups using the lm function in R. R2 represents the proportion of signaling variance explained by the predictor, and P values test the linear association between predictor and response. P values were not corrected for multiple comparisons and are interpreted as exploratory.

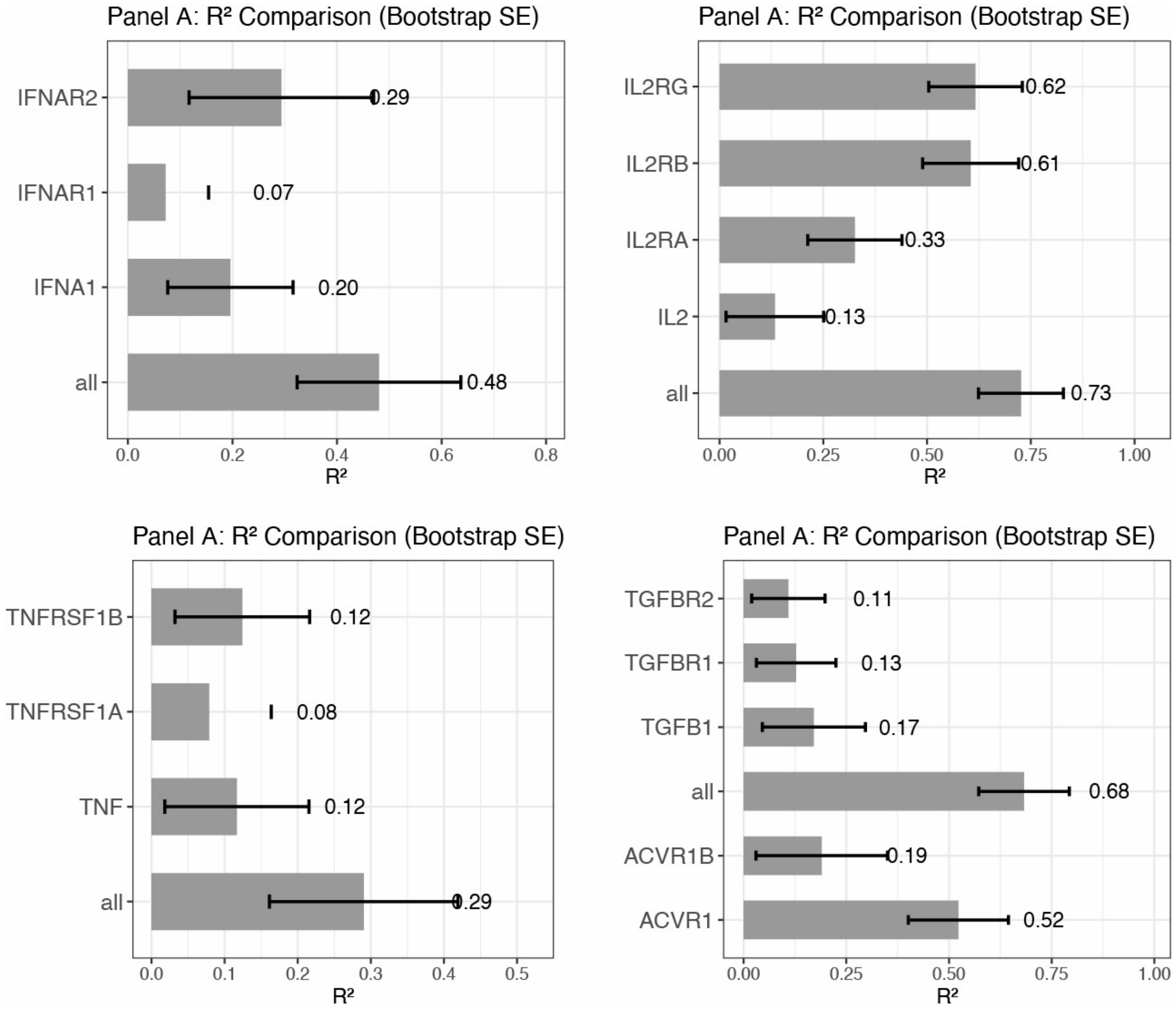

**Figure S4. Multivariable regression confirms independent contributions of multiple ligand and receptor variables to downstream signaling.** Four cytokine systems are shown in which multiple predictors exhibited nominally significant independent contributions in multivariable linear models, consistent with the central prediction of our framework. Each panel displays bootstrap $R^2$ values (1,000 iterations; error bars, 95% CI) for individual univariate predictors alongside the combined multivariable model ("all"). Top left: IFNα signaling predicted by IFNA1 ligand and IFNAR2 (combined R2 = 0.48). Top right: IL-2 signaling predicted by IL2RG, IL2RB, IL2RA, and IL2 ligand (combined $R^2$ = 0.73). Bottom left: TNF signaling predicted by TNF ligand, TNFRSF1A, and TNFRSF1B (combined R2 = 0.29). Bottom right: TGFβ signaling predicted by ACVR1, ACVR1B, TGFB1, TGFBR1, and TGFBR2 (combined R2 = 0.68). In each case, the multivariable model substantially exceeds any single predictor, confirming that ligand and receptor expression provide non-redundant information about downstream signaling in intact human tissue. _Statistical analysis_: Univariate and multivariable linear models were fit using the lm function in R, with ligand and receptor subunits entered as separate predictors. $R^2$ represents the proportion of downstream signaling variance explained by each model; 95% confidence intervals were estimated by bootstrap resampling (1,000 iterations). Individual predictor P values were obtained from coefficients in the multivariable models and were not corrected for multiple comparisons.

#### 1.3 Clinical Validation of Avidity Model Across 510 Lung Cancer Patients

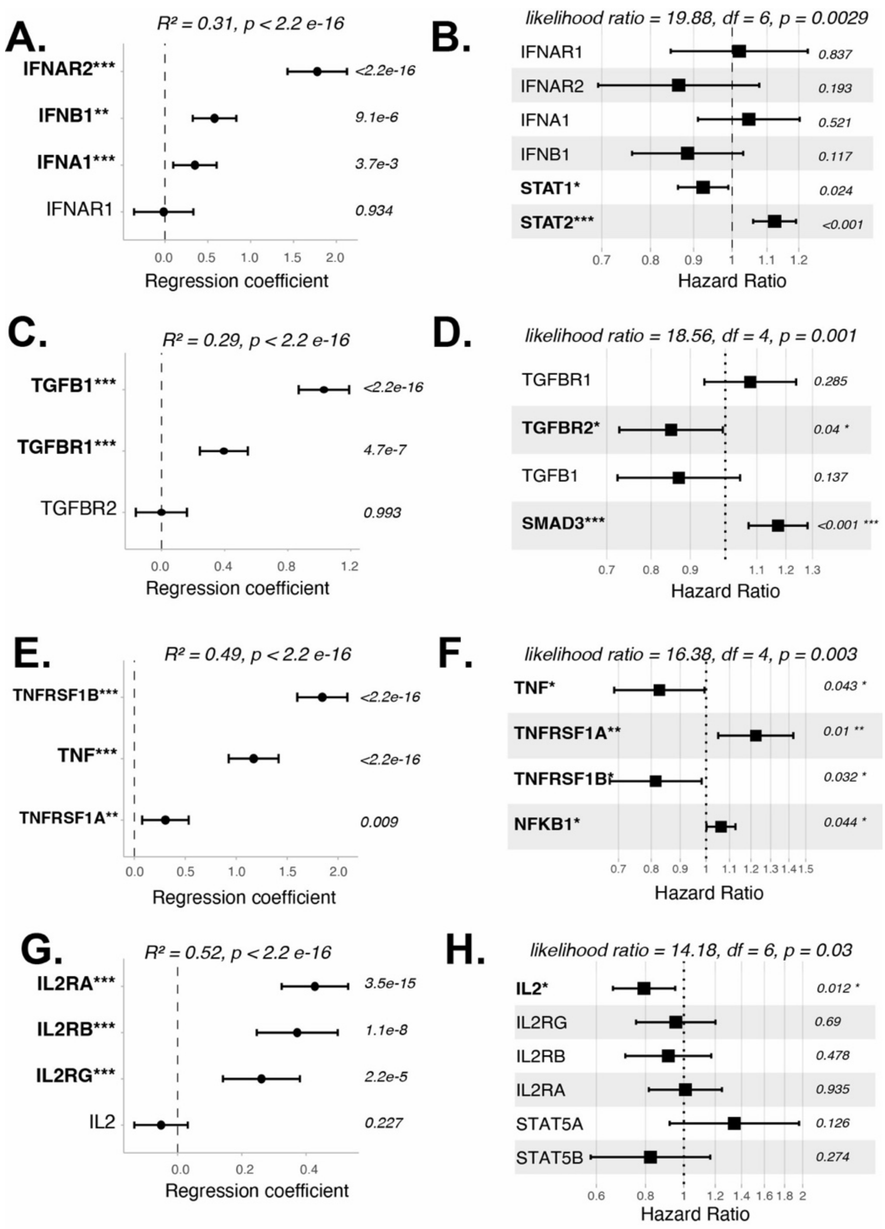

**Figure S5. Validation of cytokine signaling models across the TCGA lung adenocarcinoma cohort.** Bulk RNA-seq data from 510 patients in the TCGA Lung Adenocarcinoma (LUAD) PanCancer Atlas cohort were used to test whether cytokine signaling relationships identified by CosMx spatial transcriptomics generalize across patients. (A, C, E, G) Multivariable linear regression for IFN, TGFβ, TNF, and IL-2 signaling, respectively. Across all four cytokine systems, multiple ligand and receptor components independently predicted downstream transcription factor regulon activity, recapitulating the multivariable relationships observed in the single-cell CosMx analysis and demonstrating their conservation across patients. Points indicate regression coefficients with 95% confidence intervals; overall model $R^2$ and P values are shown above each panel. (B, D, F, H) Multivariable Cox proportional-hazards models incorporating ligand expression, receptor expression, and downstream transcription factor regulon activity for the corresponding cytokine systems. All four signaling modules were significantly associated with overall survival by likelihood-ratio testing, supporting the clinical relevance of variation within these cytokine signaling axes. Squares indicate hazard ratios with 95% confidence intervals; individual coefficient P values are shown at right. *Statistical analysis:* Panels A, C, E, and G show multivariable linear regression models fit in R, with $R^2$ representing the proportion of downstream TF-regulon variance explained by the complete ligand–receptor model; coefficient P values are from the fitted linear models. Panels B, D, F, and H show multivariable Cox proportional-hazards models; P values above each panel are from the overall likelihood-ratio test, while values beside individual terms correspond to coefficient-level tests. P values were not corrected for multiple comparisons.

#### 1.4 Clinical Validation of Avidity Models Assumptions leveraging Quantitative Proteomics of PBMCs

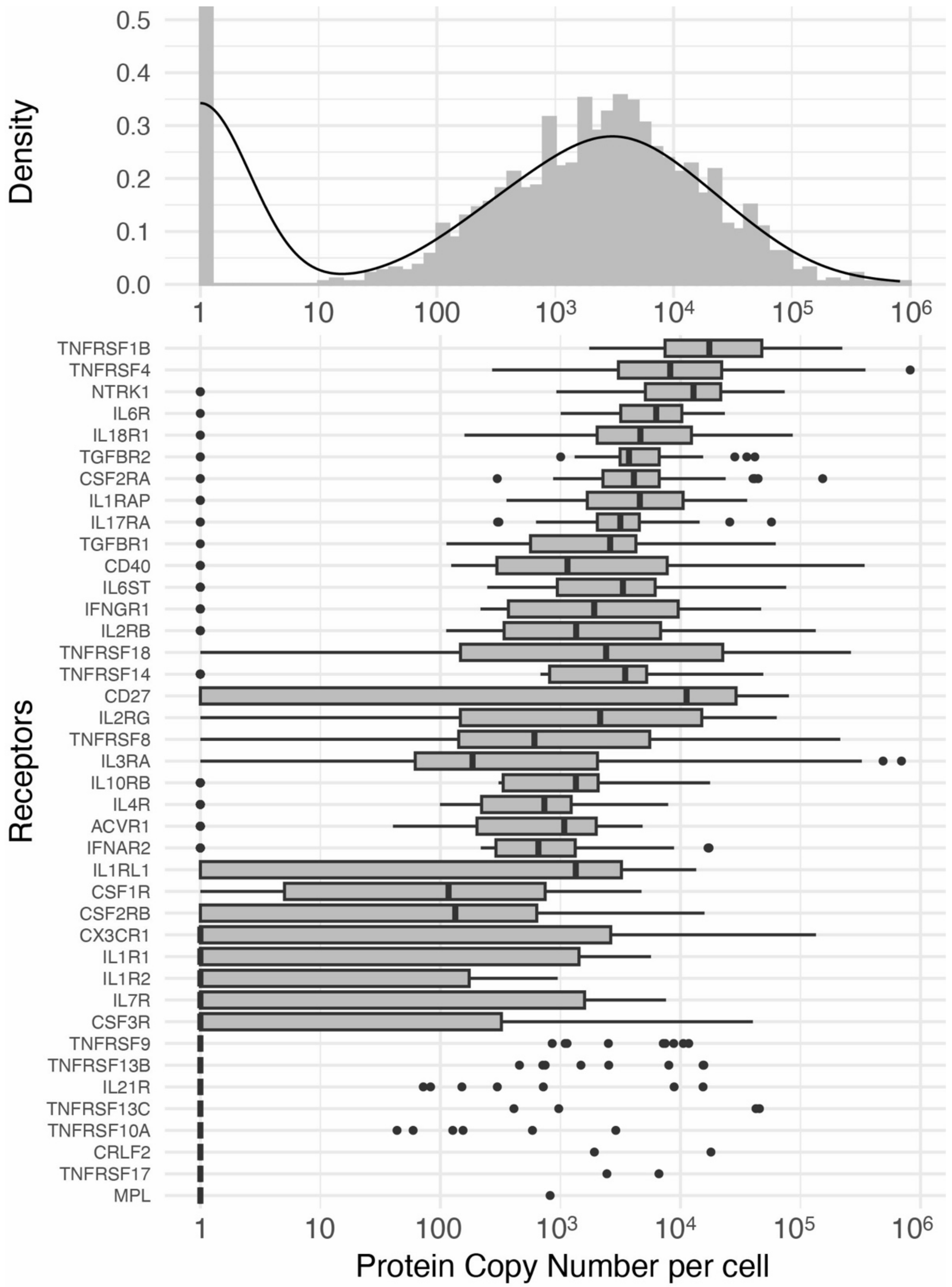

**Figure S6. Distribution of cytokine receptor protein abundance across primary human immune cells.** Quantitative proteomic measurements from Rieckmann et al. were used to estimate cytokine receptor abundance as protein copies per cell across human immune cell populations. (Top) Distribution of receptor protein copy numbers, illustrating the broad dynamic range of surface receptor abundance. (Bottom) Receptor-specific variation in protein copy number across immune cell types, shown on a logarithmic scale. Highly expressed receptors are represented by thousands to tens of thousands of copies per cell and are therefore more readily approximated by bulk concentration-based models, whereas low-abundance receptors may be increasingly sensitive to discrete or stochastic effects. These data also provide an orthogonal benchmark for transcriptomic receptor measurements, which show an overall mRNA–protein correlation of approximately $r = 0.5$ in this dataset. Boxplots show median, interquartile range, and 1.5× IQR whiskers; points indicate outliers. *Statistical analysis:* mRNA and protein abundance were $\log10(x + 1)$-transformed, and correspondence was quantified using Pearson linear correlation across matched immune-cell populations. Correlation coefficients are presented descriptively and were not corrected for multiple comparisons.

#### 1.5 Comparison of Avidity Model procedure vs common scRNAseq cell-communication methods

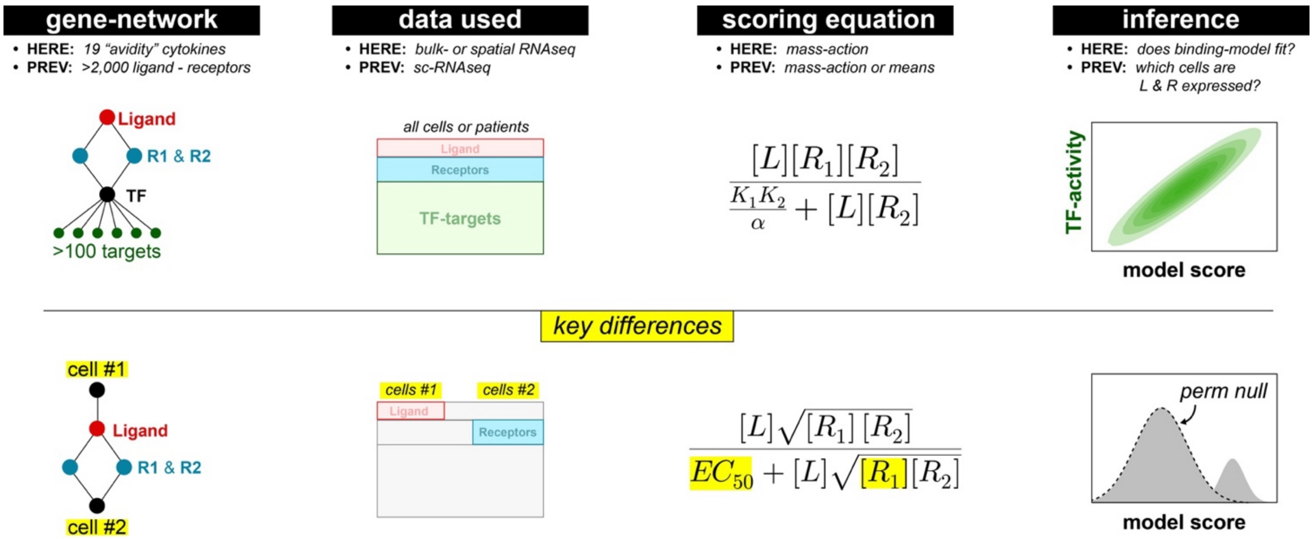

**Figure S7. Comparison of the avidity-based signaling framework with CellChat. Schematic comparison of the network representation, input data, scoring equation, and statistical inference used by the present framework (top) and CellChat (bottom).** *Top:* Our framework focuses on 19 bona fide avidity-dependent cytokine systems and jointly models ligand expression, individual receptor subunits, and downstream transcription factor regulon activity. Rather than assigning ligand and receptor expression to selected individual cells or imposing expression thresholds, the full bulk or spatial RNA-seq expression matrix is used to evaluate whether observed signaling follows the derived mass-action relationship. Model fit is assessed from the joint probability distribution between predicted signaling scores and downstream regulon activity. *Bottom:* CellChat represents ligand-receptor interactions between designated sender and receiver cells, subsets ligand and receptor expression according to cell-level expression criteria, and calculates a communication score relative to a permuted null distribution. Its Hill-type scoring equation provides the closest existing analogue to our formulation, but differs mechanistically by combining receptor subunits using a geometric mean, thereby enforcing receptor symmetry, and by using a common EC50 parameter across ligand-receptor interactions. Yellow highlighting denotes key differences from the avidity-based framework.

#### 1.6 Practical Workflow for Applying Avidity Framework to Transcriptomic Data

The avidity framework can be applied to bulk, single-cell, or spatial transcriptomic datasets when expression measurements are available for a cytokine ligand, the individual subunits of its signaling receptor complex, and a downstream transcriptional readout. The analysis does not require absolute receptor concentrations or independently measured binding constants when the goal is to test the relative ligand–receptor dependencies predicted by Equation 6a. A general workflow is outlined below. The Top of Figure S7 is repeated below to illustrate criteria for this procedure:

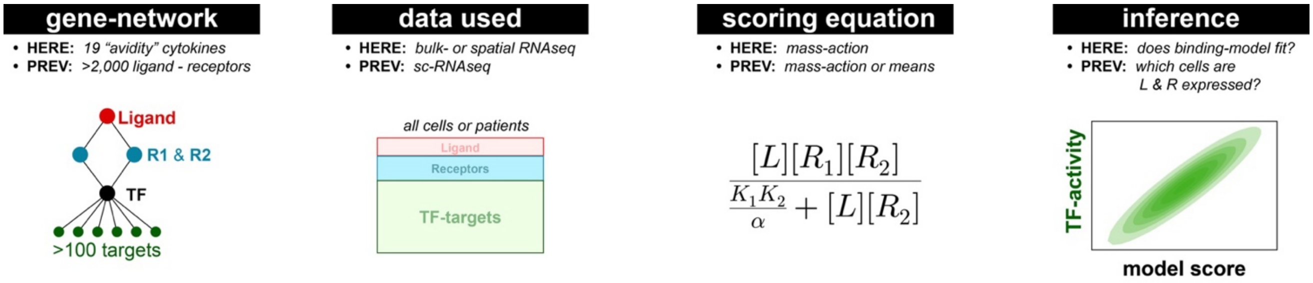

- **Step 1. Define the cytokine–receptor–response network:** Select cytokine systems for which multicomponent receptor engagement or receptor oligomerization is an established requirement for signaling.[2, 3] For each cytokine, define: (i) the ligand or ligands, (ii) each receptor subunit required for formation of the signaling complex,[2, 3] and (iii) a canonical downstream transcription factor or transcriptional signature.[4] In this study, ligand–receptor assignments were derived from CellChat and curated literature, while downstream transcription-factor relationships were defined from established signaling pathways.[4] The complete network used here contained 19 cytokine ligand-response axes.
- **Step 2. Confirm that all required signaling components are measured:** Restrict analysis to cytokine systems for which the ligand, required receptor subunits, and downstream response can all be quantified in the dataset. Missing receptor subunits should not be treated as zero expression unless their absence is biologically established; rather, the corresponding signaling axis should generally be excluded from quantitative analysis. For targeted spatial panels, this filtering may substantially reduce the number of analyzable cytokine systems.
- **Step 3. Preprocess expression measurements on a comparable scale:** Gene-expression measurements should be normalized using an approach appropriate for the experimental platform and transformed to reduce the influence of highly skewed expression distributions. Because Equation 6a becomes additive after logarithmic transformation, log-scaled expression is particularly convenient for regression-based application. In the present analyses, RNA abundance was represented using log-transformed normalized expression values. Predictor variables may additionally be standardized when direct comparison of regression coefficients across genes is desired.
- **Step 4. Define the biologically relevant ligand exposure:** Ligand expression should represent the extracellular cytokine environment experienced by the responding cell or population rather than necessarily the ligand transcripts within the responding cell itself. For bulk RNAseq, sample-level ligand expression can be used directly. For spatial transcriptomics, ligand exposure can instead be estimated from the local neighborhood surrounding each responding population. In the CosMx analysis used here, neighboring cells within a 50µm radius were summarized into local tissue niches, and cytokine expression was estimated at the niche level. Critically, this step is not possible with single-cell RNAseq.
- **Step 5. Treat receptor abundance and downstream signaling as response-cell-intrinsic variables:** Individual receptor-subunit expression should be measured within the responding cell type or population. Receptor subunits should be entered as separate predictors rather than collapsed into a geometric mean or other symmetric receptor score, because the avidity model predicts that individual receptor components can contribute asymmetrically to potency and maximal signaling. Downstream signaling should likewise be quantified within the responding population. Suitable readouts include activity of a canonical transcription-factor regulon, pathway-level transcriptional signatures, or another independently defined signaling output. In this study, we used both Hallmark pathway signatures and VIPER/DoRothEA transcription-factor regulon activity (Figure S2).

- **Step 6. Aggregate sparse single-cell measurements when necessary:** For single-cell or spatial datasets, individual-cell measurements nearly always too sparse for stable quantitative regression. Cells can therefore be aggregated into biologically meaningful pseudobulk groups while preserving the variables needed for inference. In our CosMx implementation, cells were grouped jointly by cell type and spatial neighborhood, producing cell-type x niche pseudobulk profiles. Groups with insufficient representation should be removed before regression; here, analyses were restricted to cell types constituting >5% of cells within a niche to avoid unstable estimates from rare populations.
- **Step 7. Fit the ligand-receptor-response model**: Under sub-saturating conditions, Equation 6a predicts that ligand abundance and the individual receptor components jointly contribute to signaling. The practical statistical implementation is therefore a multivariable linear model of the general form:
    - Downstream signaling ~ ligand + receptor 1 + receptor 2 + ...

  For higher-order receptor systems, all required receptor components can be included as separate predictors. Model coefficients quantify the direction and relative contribution of each signaling component, while model $R^2$ quantifies the fraction of downstream signaling variance explained by the joint ligand–receptor relationship. Univariate models can additionally be fit for comparison with the multivariable model.
- **Step 8. Interpret receptor coefficients in the context of ligand concentration and stoichiometry:** Regression coefficients should not be interpreted as fixed biochemical properties of a receptor subunit. The designation of a receptor as “limiting” or “excess” depends on its abundance relative to the other components of the signaling complex in the cellular context being analyzed. Under sub-saturating conditions, both ligand and receptor abundance can contribute to response, whereas under strongly saturating ligand conditions the model predicts that signaling becomes predominantly sensitive to the limiting receptor. Experimental knowledge of cytokine dose or receptor stoichiometry should therefore be used when assigning mechanistic interpretations to individual coefficients.
- **Step 9. Treat binding affinity and cooperativity as effective parameters unless independently measured:** Absolute prediction of $EC_{50}$ requires binding constants, receptor abundance, and cooperativity parameters on compatible physical scales. These measurements are rarely available in transcriptomic datasets. For regression-based applications, terms such as K1K2/α are therefore absorbed into the fitted intercept or effective-affinity term rather than estimated independently. Consequently, α cannot be inferred separately from affinity constants without additional biophysical measurements. The transcriptomic implementation tests the predicted dependence of signaling on ligand and receptor abundance rather than attempting to reconstruct an absolute molecular EC50.
- **Step 10. Evaluate the validity of mRNA as a receptor-abundance surrogate:** Transcript abundance should be regarded as an informative but imperfect proxy for surface receptor protein abundance. Practically, method like flow cytometry (single-cells) or immunohistochemistry (tissue) should always be used as a validation step for mRNA-based inferences. If protein measurements are unavailable, analyses should emphasize relative differences across samples or cell populations rather than interpret transcript abundance as an absolute surface concentration. In the primary human immune-cell datasets examined here, receptor mRNA and quantitative protein abundance showed an overall positive correspondence of approximately r = 0.5, while relative differences among receptor subunits were frequently preserved.
- **Step 11. Perform sensitivity and validation analyses:** Whenever possible, model relationships should be evaluated across independent datasets, signaling readouts, or experimental modalities (Figures 1-4). Appropriate validation may include replication in a separate cohorts (Figure 5), comparison of transcription-factor and pathway-level response measures (Figure S2), protein-level confirmation of receptor abundance (Figure 5D-F, S6), or perturbation experiments performed under defined cytokine concentrations (Figure 4).

In summary, the transcriptomic application of the avidity framework requires three principal measurements (ligand abundance, individual receptor-subunit abundance, and a downstream response) which it preserves the individual receptor components rather than collapsing them into a single receptor score. The resulting multivariable model provides a practical means of testing whether signaling variation follows the ligand- and receptor-dependent relationships predicted by multivalent receptor binding, while absolute EC50 prediction requires additional biophysical information.

### 2. Equilibrium Model Derivations and Underlying Assumptions

#### 2.1 The Thermodynamic Effects of Surface Immobilization on Complex formation

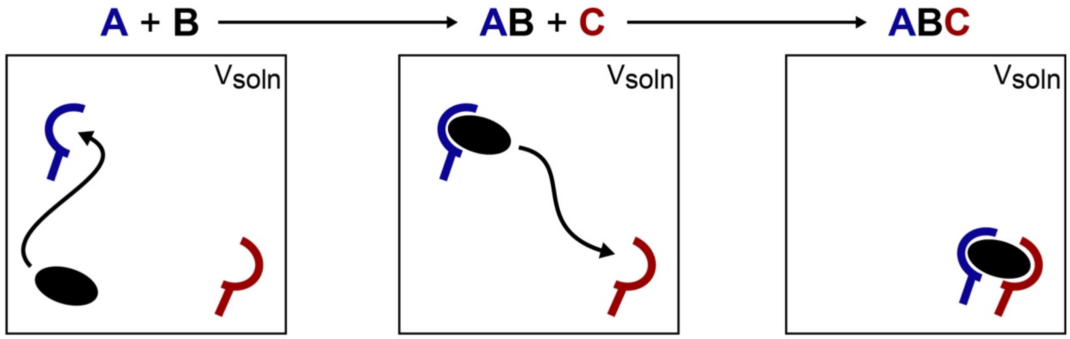

**Figure S8.** Cartoon Summary of Translational entropy changes at each stage in assemble of a bifunction ligand (B) with two receptors A and C. The search space (Vsoln) and translational entropy-change does not change appreciably with each binding event.

The predominant effect of immobilization of dimeric receptors on a cell-surface is entropic as the translational entropy of a ligand is restricted to the surface of the cell upon the first binding event (Figure S8-9). This reduction in translational entropy can be estimated by the ratio of the total solution volume ($V_{soln}$) and the surface of the cell ($V_{surf}$) by equation S1 where R is the gas constant:[5-7]

$$\Delta S = Rln\left(\frac{V_{soln}}{V_{surf}}\right) \qquad \text{S01}$$

$V_{surf}$ is typically estimated as the product of the surface area (SA) of the cell and a shell of thickness (T) around that surface that captures the approximate translational volume of surface-immobilized proteins[5, 7]

$$V_{surf} = SA \cdot T \qquad \text{S02}$$

A common estimate for T is 10nm which has been often used for the type-1 interferons.[8-11] In this work, this assumption is not used as we do not attempt to directly parameterize our models and instead fit for parameters empirically using the form of our analytical solutions.

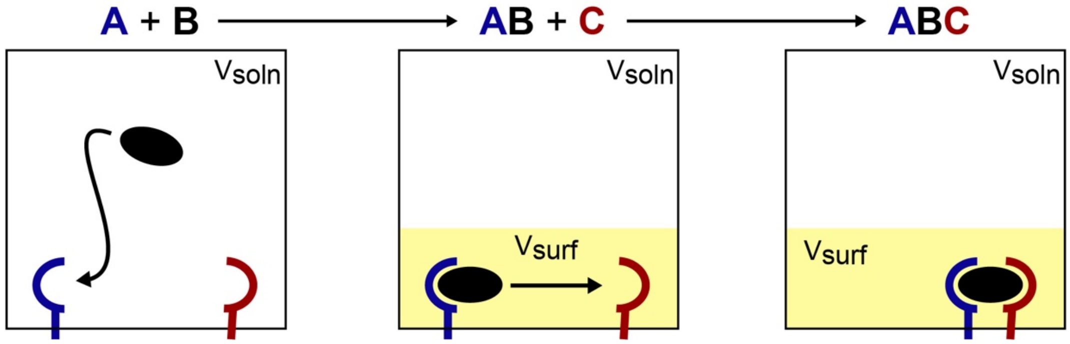

**Figure S9** Cartoon Summary of Translational entropy changes at each stage in assemble of a bifunction ligand (B) with two receptors A and C that are immobilized on a cell-surface. The search space ($V_{soln}$) and translational entropy-change for the second binding event is reduced as a result of the increased "effective concentration" of the immobilized ligand.

Critically, the first binding increases the affinity of the second binding event by reducing the entropic penalty of binding the second receptor. This phenomenon has been referred to "cell-surface cooperativity" and an enhanced "effective concentration" of immobilized ligand.[5-7, 12] Practically, this entropic boost in affinity can be thought of as a reduction in the search-space needed to find the second receptor.

#### 2.2 Derivations of Equilibrium Models of Multi-Receptor Avidity

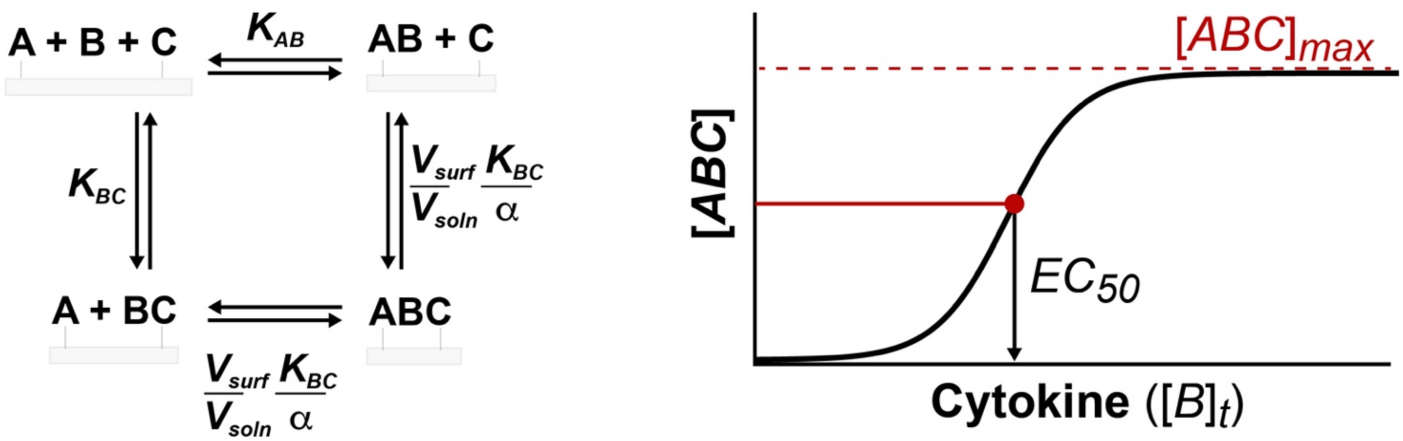

**Figure S10.** Equilibrium scheme (right) and $EC_{50}$ calculation procedure detailed below. Equilibrium scheme: Assembly of the ABC complex can occur via two paths where the initial binding of A to B or B to C is described by an equilibrium binding constants KAB and KBC respectively. Assembly of complete ABC complex must account for non-binding interactions between receptors A and B ($\alpha$) and the "effective concentration" enhancement described in the previous section. *EC50 Calculation Procedure*: $[ABC]_{max}$ is first estimated and $[ABC]_{max}/2$ is used to calculate the EC50 using a closed-form solution for total bridging cytokine B.

Figure S10's scheme describes the various paths available for the binding of a bridging ligand (B) to two cell-surface receptors: A and C. "A" and "C" are used here instead of "R1" and "R2" (used in the main paper) to maintain clarity in the derivations below and maintain derivation consistency with our previous work where several details proofs and sensitivity analyses are presented.[13] "A" is equivalent to "R1", and "C" is equivalent to "R2" throughout.

##### *Note on Cooperativity Conventions (antibodies and cytokins)*

**Cooperativity Conventions:** This use of α follows the conventional thermodynamic treatment of ternary-complex binding, in which formation of one binary interaction creates a new molecular species whose affinity for the subsequent binding partner may differ from that of the isolated components.[14] Operationally, α scales the affinity of the subsequent binding event: $\alpha > 1$ represents positive cooperativity or stabilization of the ternary complex, $\alpha = 1$ represents neutral or independent binding, and $\alpha < 1$ represents negative cooperativity or destabilization.[14, 15] In classical allosteric ternary-complex models, for example, the dissociation constant for the second binding event becomes $Kd/\alpha$, with thermodynamic reversibility requiring the same coupling factor regardless of the order in which the complex is assembled. Thus, α should be interpreted as a thermodynamic coupling term rather than as an additional binding site or receptor concentration.

**Cytokine Cooperativity**: Such coupling has long been recognized in cytokine-receptor assembly. Early equilibrium models of aggregating cytokine receptors explicitly allowed ligand binding and receptor association to alter one another's apparent affinities.[16] Whitty *et al.* subsequently quantified this coupling experimentally for heteromeric cytokine receptors, showing that recruitment of the common γ-chain stabilized ligand-bound IL-4 and IL-2 receptor complexes, with dimensionless interaction factors of approximately 9-10 under the cellular conditions examined.[17, 18] Importantly, cooperativity is system-specific rather than universally positive: membrane-reconstituted measurements of the IFNα2–IFNAR1–IFNAR2 ternary complex found no detectable additional cooperative effect beyond the constituent binding interactions.[19] Accordingly, α is retained in the general equations to accommodate cytokine-specific coupling, while $\alpha = 1$ corresponds to the non-cooperative limit.

**Antibody Cooperativity:** For the antibody systems considered here, we assume $\alpha = 1$, corresponding to no additional allosteric coupling between the two Fab antigen-binding events. This assumption is supported by the high structural flexibility of intact IgG, in which the hinge permits the two Fab arms to adopt widely different orientations and move largely independently.[20] Structural studies likewise show that the two antigen-binding sites remain highly similar despite substantial differences in Fab orientation.[20] More broadly, IgG avidity is generally attributed to associative crosslinking rather than antigen-induced intramolecular allostery.[21] Accordingly, $\alpha = 1$ provides an appropriate minimal assumption for the antibody avidity calculations used here.

***i. Heterodimeric Receptor Complexes on a Cellular Surface***

Figure S10's equilibrium scheme can be defined mathematically using the *conservation of mass* which restricts the concentrations of free species ([A], [B], [C], [AB], [BC], [ABC]) by the total concentration of A, B and C ($[A]_t$, $[B]_t$, $[C]_t$):

$$[A]_t = [A] + [AB] + [ABC] \qquad \text{S03}$$

$$[B]_t = [B] + [AB] + [BC] + [ABC] \qquad \text{S04}$$

$$[C]_t = [C] + [BC] + [ABC] \qquad \text{S05}$$

And the *law of mass action* which defines the favorability of binding events via the dissociation constants $K_{AB}$ and $K_{BC}$:

$$K_{AB} = \frac{[A][B]}{[AB]} \qquad \text{S06}$$

$$K_{BC} = \frac{[B][C]}{[BC]} \qquad \text{S07}$$

For the ternary complex assembly, the energetic favorability is captured by both constituent binding events, a cooperativity factor ($\alpha$) which captures stabilizing or destabilizing interactions between A and C (which aren't directly bound) and the entropic-cell-surface cooperativity factor described in the previous section.

$$\frac{K_{AB}K_{BC}}{\alpha}\frac{V_{surf}}{V_{soln}} = \frac{[A][B][C]}{[ABC]} \qquad \text{S08}$$

For simplicity, as $\alpha$ and the volume ratio (Figure S5) have the same effect we combine them into one term $\alpha_{surf}$ $(= \alpha \cdot \frac{V_{soln}}{V_{surf}})$

$$\frac{K_{AB}K_{BC}}{\alpha_{surf}} = \frac{[A][B][C]}{[ABC]} \qquad \text{S09}$$

Previously we have shown that the above system of 6 equations and 6 variables ([A], [B], [C], [AB], [BC], [ABC]) can be solved for [ABC]. Yielding the below quintic equation:[13]

$$\begin{aligned}
&[\boldsymbol{ABC}]^{\mathbf{5}}(\alpha-1)\alpha^2 - \\
&[\boldsymbol{ABC}]^{\mathbf{4}}(\alpha^2([A]_t+[B]_t+2[C]_t)+2\alpha(K_{AB}+K_{BC}-[A]_t-[C]_t)-2(K_{AB}+K_{BC}))+ \\
&[\boldsymbol{ABC}]^{\mathbf{3}}\left(\begin{matrix}\alpha^3(([A]_t)^2+2[A]_t([B]_t+2[C]_t)+[C]_t(2[B]_t+[C]_t))+ \\ \alpha^2\left(\begin{matrix}-[A]_t([B]_t+3[C]_t-2K_{AB}-3K_{BC})+K_{BC}([B]_t+2[C]_t+K_{AB})- \\ ([A]_t)^2+([B]_t)^2-[B]_t[C]_t-([C]_t)^2+[B]_tK_{AB}+3[C]_tK_{AB}\end{matrix}\right)+ \\ \alpha\big((K_{BC})^2-2K_{BC}([A]_t+[C]_t+K_{AB})+K_{AB}(K_{AB}-2([A]_t+[C]_t))\big)- \\ (K_{AB}+K_{BC})^2\end{matrix}\right)+ \\
&[\boldsymbol{ABC}]^{\mathbf{2}}\boldsymbol{\alpha}\left(\begin{matrix}\alpha^2([B]_t([C]_t)^2+[A]_t[C]_t(2[B]_t+[C]_t)+([A]_t)^2\,([B]_t+2[C]_t))- \\ \alpha\left(\begin{matrix}-([A]_t)^2([B]_t+[C]_t-K_{BC})+ \\ [A]_t(([B]_t)^2+[B]_t(K_{AB}+K_{BC}-2[C]_t)-([C]_t)^2+3[C]_t(K_{AB}+K_{BC})+K_{AB}K_{BC})+ \\ [C]_t\big(([B]_t)^2+[B]_t(K_{AB}+K_{BC}-[C]_t)+K_{AB}([C]_t+K_{BC})\big)\end{matrix}\right)\end{matrix}\right)+
\end{aligned} \qquad \text{S10}$$

$$[\boldsymbol{ABC}]\alpha^2[A]_t[C]_t\begin{pmatrix}([B]_t)^2+[B]_t(2\alpha[C]_t+K_{AB}+K_{BC}-[C]_t)+[A]_t\big((2\alpha-1)[B]_t+\alpha[C]_t+K_{BC}\big)\\+K_{AB}([C]_t+K_{BC})\end{pmatrix}-$$
$$\alpha^3([A]_t)^2\,[B]_t([C]_t)^2=0$$

Unfortunately, we have also demonstrated that this 5th order polynomial is algebraically unsolvable except for the maximum value $[ABC]_{max}$.[13] In this current work, we are not interested in [ABC] per se but rather the $EC_{50}$ for ABC-response curves which corresponds to the concentration of B ($[B]_t$) at which [ABC] = $[ABC]_{max}$ / 2.

Rearranging equation S10, to solve for [B]t yields a 2nd order polynomial which can be solved with the quadratic formula:

$$[B]_t = \left(\frac{[A]_t+[C]_t-K_{AB}-K_{BC}+\frac{\alpha_{surf}([ABC]-[A]_t)\,([ABC]-[C]_t)}{[ABC]}+\frac{[A]_t(K_{BC}-K_{AB})}{\alpha_{surf}([ABC]-[A]_t)}+\frac{[C]_t(K_{AB}-K_{BC})}{\alpha_{surf}([ABC]-[C]_t)}\pm\left(\frac{\alpha_{surf}}{[ABC]}+\frac{1}{[A]_t-[ABC]}+\frac{1}{[C]_t-[ABC]}\right)\sqrt{\begin{matrix}\big(\alpha_{surf}([ABC]-[A]_t)([ABC]-[C]_t)-[ABC]K_{AB}\big)^2-\\2[ABC]K_{BC}\big(\alpha_{surf}([ABC]-[A]_t)([ABC]-[C]_t)+[ABC]K_{AB}\big)+\\{[ABC]}^2{K_{BC}}^2\end{matrix}}}{\alpha_{surf}}\right) \quad \text{S11}$$

Previously we have derived a general solution for [ABC]max

$$[ABC]_{max}=\frac{[A]_t+[C]_t+\frac{\left(\sqrt{K_{AB}}+\sqrt{K_{BC}}\right)^2}{\alpha_{surf}}-\sqrt{\left([A]_t+[C]_t+\frac{\left(\sqrt{K_{AB}}+\sqrt{K_{BC}}\right)^2}{\alpha_{surf}}\right)^2-4[A]_t[C]_t}}{2} \quad \text{S12}$$

Assuming that A is the limiting reagent, we have previously shown that the necessary cooperativity for $[ABC]_{max}$ to occur at half-the limiting reagent (A) is: [13]

$$\alpha_{crit}=\frac{\left(\sqrt{K_{AB}}+\sqrt{K_{BC}}\right)^2}{[C]_t-[A]_t/2} \quad \text{S13}$$

Under high cooperativity systems ($\alpha>\alpha_{crit}$) – expected for cell-surface receptors where signaling complex forms appreciably – equation S12 can be approximated by:

$$[ABC]_{max}=\frac{[A]_t}{2} \quad \text{S14}$$

Plugging S14 in for [ABC]max in equation S11 results in extensive cancelation of terms, most notably removal of the root term. Additional assumption of quadrant 1 conditions in our previous work (full derivation can be found in reference 10),[13] that are typical for receptor-bound ternary systems, results in the final equation for EC50:

$$EC_{50}=\frac{[A]_t}{2}+\frac{K_{AB}K_{BC}}{\alpha_{surf}([C]_t-[A]_t/2)} \quad \text{S15}$$

Noting that the solution concentration ($V_{soln}$) is implicit in concentrations of $[C]_t$ and $[A]_t$. (moles per liter) we can remove the Volume terms from $\alpha_{surf}$:

$$EC_{50} = \frac{[A]_t}{2} + \frac{K_{AB}K_{BC}}{\alpha \frac{V_{soln}}{V_{surf}}([C]_t - [A]_t/2)} \quad \text{S16}$$

And convert some of the concentration terms to "surface molarity" concentrations (Figure S4):

$$EC_{50} = \frac{[A]_t}{2} + \frac{K_{AB}K_{BC}}{\alpha\left([C]_t^{surf} - [A]_t^{surf}/2\right)} \quad \text{S17}$$

Noting that the value of $\left([C]_t^{surf} - [A]_t^{surf}/2\right)$ can only range between $[C]_t^{surf}$ and $[C]_t^{surf}/2$ and cell-surface receptors correspond to quadrant 1 ($[A]_t \ll K_{AB}$, $[C]_t \ll K_{BC}$,) in our previous publication[13] we can obtain the following simple equation to estimate EC50:

$$EC_{50} = \frac{K_{AB}K_{BC}}{\alpha[C]_t^{surf}} \quad \text{S18}$$

***ii. Homodimeric Receptor Complexes on a Cellular Surface***

The equilibrium scheme for ligands (B) which homodimerize the same receptor(A) can be defined mathematically using the conservation of mass which restricts the concentrations of free species ([A], [B], [C], [AB], [ABA]) by the total concentration of A and B ($[A]_t$, $[B]_t$):[15]

$$[A]_t = [A] + [AB] + 2[ABA] \quad \text{S19}$$

$$[B]_t = [B] + [AB] + [ABA] \quad \text{S20}$$

And the law of mass action which defines the favorability of binding events via the dissociation constants $K_{AB}$ and $K_{BC}$:

$$\frac{K_{AB}}{2} = \frac{[A][B]}{[AB]} \quad \text{S21}$$

For the ternary complex assembly, the energetic favorability is captured by both constituent binding events, a cooperativity factor ($\alpha$) which captures stabilizing or destabilizing interactions between A and C (which aren't directly bound) and the entropic-cell-surface cooperativity factor described in the previous section

$$\frac{2K_{AB}}{\alpha}\frac{V_{surf}}{V_{soln}} = \frac{[AB][A]}{[ABA]} \quad \text{S22}$$

Notabily the "2" terms in equations S21 and S22 are derived from statistical factors accounting for the symmetric nature of B were on initial binding is 2x favorable as two A-B binding sites are available upon initial binding relative to the total number of B molecules. Whitesides and coworkers describe this factor in more detail in their original derivation.[15] For simplicity, as $\alpha$ and the volume ratio have the same effect we combine them into one term $\alpha_{surf}$

$$\frac{2K_{AB}}{\alpha_{surf}} = \frac{[AB][A]}{[ABA]} \quad \text{S23}$$

The above system of equation has already been previously solved for the solution case which we here extend to cell-surface dimerization. First [ABA]max has been previously derived as:[15]

$$[ABA]_{max} = \frac{[A]_t + \frac{2K_{AB}}{\alpha} - \sqrt{4\frac{K_{AB}}{\alpha}\left([A]_t + \frac{K_{AB}}{\alpha}\right)}}{2} \quad \text{S24}$$

Noting that the critical cooperativity need to elicit a half-maximal $[ABA]_{max}$ must be a multiple of $K_{AB}/[A]_t$ we replaced $[ABA]_{max}$ with $[A]_t/4$ and a with $xK_1/[A]_t$ and solved for x:

$$\frac{[A]_t}{4} = \frac{[A]_t + \frac{2[A]_t}{x} - \sqrt{4\frac{[A]_t}{x}\left([A]_t + \frac{[A]_t}{x}\right)}}{2} \quad \text{S25}$$

$$x = 8 \quad \text{S26}$$

$$\alpha_{crit} = 8\frac{K_{AB}}{[A]_t} \quad \text{S27}$$

Under high cooperativity systems ($\alpha > \alpha_{crit}$) – expected for cell-surface receptors where signaling complex forms appreciably – quatitative ternary complex forms:

$$[ABA]_{max} = \frac{[A]_t}{2} \tag{S28}$$

As a result minimal [AB] forms when $[B]_t$ < [B]t,max and can be removed from system of equations S19-21

$$[A]_t = [A] + 2[ABA] \tag{S29}$$

$$[B]_t = [B] + [ABA] \tag{S30}$$

$$\frac{(K_{AB})^2}{\alpha_{surf}} = \frac{[A][B][A]}{[AB]} \tag{S31}$$

Eliminating [A] and [B] and solving for [B]t:

$$[B]_t = \frac{4\alpha_{surf}[ABA]^3 - 4\alpha_{surf}[ABA]^2[A]_t + 4\alpha_{surf}[ABA][A]_t{}^2 + [ABA]K_{AB}{}^2}{\alpha_{surf}(2[ABA] - [A]_t)^2} \tag{S32}$$

Replacing [ABA] with [A]t/4 and simplifying we obtain the following equation for $EC_{50}$

$$EC_{50} = \frac{[A]_t}{4} + \frac{K_{AB}{}^2}{\alpha_{surf}[A]_t} \tag{S33}$$

Converting surface cooperativity to surface concentration term as we did for ABC:

$$EC_{50} = \frac{[A]_t}{4} + \frac{K_{AB}{}^2}{\alpha[A]_t^{surf}} \tag{S34}$$

Noting that for cell-surface systems the total solution concentration of receptors would be significantly restricted $[A]_t \ll K_{AB}$, equation S34 can be simplified to:

$$EC_{50} = \frac{K_{AB}{}^2}{\alpha[A]_t^{surf}} \tag{S35}$$

***iii. Homotrimerization of Cell-surface Receptors (Tumor Necrosis Factor)***

For homo-trimerizing cell-surface receptors the Conservation of Mass and Law of Mass Action Equations are as follows: [15]

$$[A]_t = [A] + [AB] + 2[ABA] + 3[A_3B] \quad \text{S36}$$

$$[B]_t = [B] + [AB] + [ABA] + [A_3B] \quad \text{S37}$$

$$\frac{K_{AB}}{3} = \frac{[A][B]}{[AB]} \quad \text{S38}$$

$$\frac{K_{AB}}{\alpha}\frac{V_{surf}}{V_{soln}} = \frac{[AB][A]}{[ABA]} \quad \text{S39}$$

$$\frac{3K_{AB}}{\alpha}\frac{V_{surf}}{V_{soln}} = \frac{[ABA][A]}{[A_3B]} \quad \text{S40}$$

Under conditions of quantitative $A_3B$ complex formation [AB] and [ABA] should not form appreciably when $[B]_t < [B]_{t,max}$ [13] and can be removed from the system of equations S36-S40:

$$[A]_t = [A] + 3[A_3B] \quad \text{S41}$$

$$[B]_t = [B] + [A_3B] \quad \text{S42}$$

$$\frac{(K_{AB})^3}{\alpha^2}\left(\frac{V_{surf}}{V_{soln}}\right)^2 = \frac{[ABA][A]}{[A_3B]} \quad \text{S43}$$

Eliminating [A] and [B] and replacing $[A_3B]$ with $[A]_t/6$ and solving for $[B]_t$ we obtain the $EC_{50}$:

$$EC_{50} = \frac{[A]_t}{6} + \frac{4{K_{AB}}^3}{3\left(\alpha[A]_t^{surf}\right)^2} \quad \text{S44}$$

Noting that for cell-surface systems the total solution concentration of receptors would be significantly restricted $[A]_t \ll K_{AB}$, equation S44 can be simplified to:

$$\boldsymbol{EC_{50} = \frac{4{K_{AB}}^3}{3\left(\alpha[A]_t^{surf}\right)^2}} \quad \text{S45}$$

***iv. Homo-n-merization (poly-valent cellular binding e.g. IgM)***

Extending homo-trimerization model to homo-n-merization models for generic cellular we can assume quantitative n-mer complex relative to constituent complexes.[22]

$$[A]_t = [A] + n[A_nB] \quad \text{S46}$$

$$[B]_t = [B] + [A_nB] \quad \text{S47}$$

$$\frac{K_{AB}}{n} = \frac{[A][B]}{[AB]} \quad \text{S48}$$

$$\frac{K_{AB}}{\alpha}\frac{V_{surf}}{V_{soln}} = \frac{[AB][A]}{[ABA]} \quad \text{S49}$$

$$K_{AB}\left(\frac{K_{AB}}{\alpha}\frac{V_{surf}}{V_{soln}}\right)^{n-1} = \frac{[A]^n[B]}{[A_nB]} \quad \text{S50}$$

Eliminating [A] and [B], replacing $[A_nB]$ with $[A]_t/2n$ and solving for $[B]_t$ we obtained an expression for the $EC_{50}$:

$$EC_{50} = \frac{[A]_t}{2n} + K_{AB}\frac{1}{n}\left(\frac{2K_{AB}}{\alpha[A]_t^{surf}}\right)^{n-1} \quad \text{S51}$$

Noting that for cell-surface systems the total solution concentration of receptors would be significantly restricted $[A]_t \ll K_{AB}$, equation S51 can be simplified to:

$$\boldsymbol{EC_{50} = K_{AB}\frac{1}{n}\left(\frac{2K_{AB}}{\alpha[A]_t^{surf}}\right)^{n-1}} \quad \text{S52}$$

*v. Hetero-Trimerizing Cell-surface receptors (e.g. Interleukin-2)*

For hetero trimerizing cell-surface receptors The Conservation of Mass and Law of Mass Action Equations are as follows:

$$[A]_t = [A] + [AB] + [ABC] + [ABD] + [ABCD] \tag{S53}$$

$$[B]_t = [B] + [AB] + [BC] + [BD] + [ABC] + [ABD] + [BCD] + [ABCD] \tag{S54}$$

$$[C]_t = [C] + [BC] + [ABC] + [BCD] + [ABCD] \tag{S55}$$

$$[D]_t = [D] + [BD] + [ABD] + [BCD] + [ABCD] \tag{S56}$$

$$K_{AB} = \frac{[A][B]}{[AB]} \tag{S57}$$

$$K_{BC} = \frac{[B][C]}{[BC]} \tag{S58}$$

$$K_{BD} = \frac{[B][D]}{[BD]} \tag{S59}$$

$$\frac{K_{AB}K_{BC}}{\alpha}\frac{V_{surf}}{V_{soln}} = \frac{[A][B][C]}{[ABC]} \tag{S60}$$

$$\frac{K_{BC}K_{BD}}{\beta}\frac{V_{surf}}{V_{soln}} = \frac{[C][B][D]}{[CBC]} \tag{S61}$$

$$\frac{K_{AB}K_{BD}}{\gamma}\frac{V_{surf}}{V_{soln}} = \frac{[A][B][D]}{[ABD]} \tag{S62}$$

$$\frac{K_{AB}K_{BC}K_{BD}}{\alpha\beta\gamma}\left(\frac{V_{surf}}{V_{soln}}\right)^2 = \frac{[A][B][C][D]}{[ABCD]} \tag{S63}$$

Under conditions of quantitative ABCD complex formation no binary or ternary complexes should form appreciably when $[B]_t < [B]_{t,max}$ (as then they would begin to out-compete ABCD preventing it from forming quantitative complex [13]) as such these terms can be removed from the system of equations S53-S63:

$$[A]_t = [A] + [ABCD] \tag{S64}$$

$$[B]_t = [B] + [ABCD] \tag{S65}$$

$$[C]_t = [C] + [ABCD] \tag{S66}$$

$$[D]_t = [D] + [ABCD] \tag{S67}$$

$$\frac{K_{AB}K_{BC}K_{BD}}{\alpha\beta\gamma}\left(\frac{V_{surf}}{V_{soln}}\right)^2 = \frac{[A][B][C][D]}{[ABCD]} \quad \text{S68}$$

Eliminating [A], [B], [C] and [D] and replacing [ABCD] with $[A]_t/2$ and solving for $[B]_t$ we obtain the $EC_{50}$:

$$EC_{50} = \frac{[A]_t}{2} + \frac{K_{AB}K_{BC}K_{BD}}{\alpha\beta\gamma[C]_t^{surf}[D]_t^{surf}} \quad \text{S69}$$

### 3. Steady-State Model Derivations and Underlying Assumptions

#### 3.1 Kinetic Limit on Signaling Potency

##### *A. Pre-Equilibrium vs Steady State Conditions: 2-Component Michaelis Menton-type Systems*

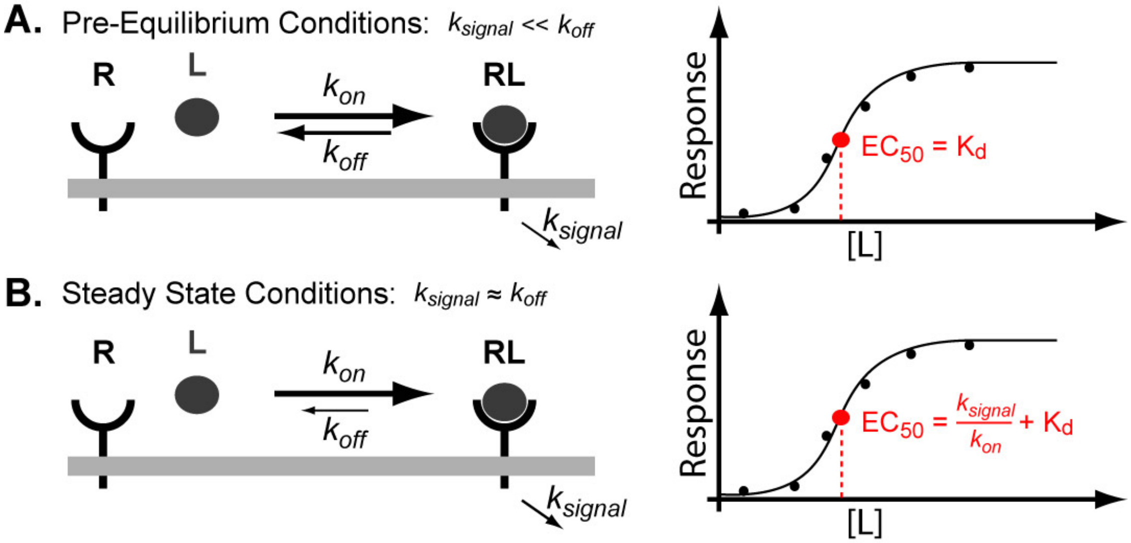

**Fig. S11 A.** Pre-equilibrium models assume that receptor ligand equilibration is much faster than signaling **B.** Steady-state models assume that receptor ligand equilibration is on approximately the same time scale as signaling

All previous derivations have relied upon the assumption of "pre-equilibrium" which asserts that the time scales of assembly and disassembly of the functional complex (RL) is faster than signaling or any other intracellular processes which affect the biological response (Figure S11A).

While this assumption is often reasonable, it breaks down when the time-scale of disassembly of the complex (RL) approaches the timescale of intracellular processes such as signaling ($k_{signal}$) or negative feedback mechanisms such as ligand-induced receptor endocytosis ($k_{endo}$).[5, 23-25]

Under such circumstances the steady-state assumption must be employed which explicitly includes contributions of additional downstream processes (Figure S11B). The steady state approximation assumes that over the timescale of the biological response the overall rates of formation and loss of the functionally relevant complex (RL) become balanced.

Both the pre-equilibrium and steady state approximations are best understood in the context of binary (2-component) Substrate/Enzyme or Ligand/Receptor interactions where the Michaelis-Menton Equation can be used (Figure S11):[25]

$$v = \frac{v_{max}[L]}{K_m + [L]} \qquad \text{S70}$$

Under pre-equilibrium conditions the potency of a ligand ($EC_{50}$) is equivalent to the binding constant for the Receptor/ligand complex because $K_m = K_d$:

$$EC_{50} = K_m = \frac{k_{off}}{k_{on}} = K_d \qquad \text{S71}$$

Under steady-state conditions the potency of a ligand ($EC_{50}$) is still equal to the Km but no longer equivalent to the binding constant of the ligand:

$$EC_{50} = K_m = \frac{k_{sig} + k_{off}}{k_{on}} = \frac{k_{sig}}{k_{on}} + K_d \qquad \text{S72}$$

Practically, it has been shown that, steady state conditions lead to the formation of a “kinetic limit” on the potency of a ligand-induced EC50’s for example, affinity maturation of antibodies exhibits a kinetic floor $EC_{50}$ ~ 10 pM.[26] This value has been explained in terms of typical values for signaling an/or endocytosis times scales and on-rates (typically invariant for protein-protein and small molecule-protein interactions):[5, 27]

$$\text{"Kinetic Limit"} = \frac{k_{sig}}{k_{on}} \approx \frac{10^{-4} s^{-1}}{10^{+6} s^{-1} M^{-1}} \approx 10\ pM \quad \text{S73}$$

Beyond this limit improvements in the binding affinity of a receptor/limit complex ($K_d$) lead to no further increase in potency ($EC_{50}$).

Below we derive $EC_{50}$ equations using the steady state approximating analogous to those derived previously. In general, they exhibit the same form as the pre-equilibrium equations with the addition of a similar “kinetic limit” term (analogous to the steady-state solution to the Michaelis-Menten kinetics).

#### 3.2 Derivations of Steady-State Models of Multi-Receptor Avidity

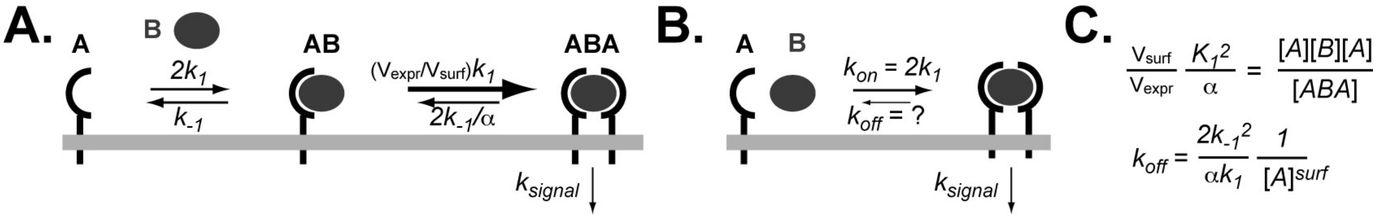

**Fig. S12 A.** Complete scheme for ABA complex formation on a cell surface **B.** Appreviated scheme for ABA complex formation on a cell surface which is accurate when ABA >> AB **C.** Derivation of $k_{off}$ when ABA >> AB on a cellular surface **D-E.** With $k_{off}$ in hand steady-state models can be easily derived with include down-stream kinetic effects

In order solve a steady-state model for multi-component (>2) complex assembly it is important to understand the rates of assembly and disassembly of that functional complex (ABA in Figure S12A). Breaking up the symmetric ABA equilibrium scheme into its constituent kinetic steps (Figure S12A) one can see that there is no single molecule step which corresponds to "assembly" and "disassembly" as there is in 2-component systems (Figure S12). As such the "timescales" of ABA assembly and disassembly are not immediately apparent but can be derived by careful consideration of the characteristics of ABA equilibria and sequential kinetic mechanisms.[28]

First, under conditions of quantitative ABA complex formation (as defined in our previous publication[13] and discussed in the previous section) ABA >> AB when $[B]_t < [B]_{t,max}$ and multi-step assembly can be approximated by the rate-determining step (first step) as illustrated in Figure S12B. Because the kinetic path does not affect the thermodynamic state changes, the governing equilibrium expression can be broken down into its constituent rate constants:

$$\frac{V_{surf}}{V_{soln}}\frac{K_1^2}{\alpha} = \frac{V_{surf}}{V_{soln}}\frac{k_{-1}^2}{\alpha k_1^2} = \frac{[A][B][A]}{[ABA]} = \frac{"k_{off}"}{"k_{on}"} \quad \text{S74}$$

While this equation does not directly give kinetic information, it provides a ratio of effective on and off rate constants for the formation of the complex ABA (under conditions where minimal AB is present at steady state) Now we can define "$k_{on}$" because we know that the rate-determining step (RDS) in ABA assembly is the formation of AB.[28] This allows us to define $k_{on}$ in Figure S12B as $2k_1$ Replacing $k_{on}$ in equation S74 with $2k_1$ and solving for $k_{off}$ we obtain:

$$k_{off} = \frac{2k_{-1}^2}{\alpha k_1}\frac{1}{[A]^{surf}} \quad \text{S75}$$

The form of this equation makes sense when one considers that disassembly of ABA requires two steps to go to completion both of which can be directly seen when separating S75 into two terms:

$$k_{off} = \frac{2k_{-1}}{\alpha k_1} \times \frac{k_{-1}}{[A]^{surf}} \quad \text{S76}$$

The first term in S76 represents the first step of disassembly ABA $\rightarrow$ AB which has the rate constant $2k_{-1}/\alpha$.[15] Once AB forms however it has a choice it can form ABA or B: ABA $\leftarrow$AB $\rightarrow$B. The rates for these respective processes are $k_1$ [A][AB] and $k_{-1}$ [AB] and the second term in S76 represents the ratio of these rates and thus the competition inherent in the second step.

##### *i. Symmetric ABA Complex EC$_{50}$*

With effective on and off rate constants defined we can now begin to model additional effects such as downstream signaling or endocytosis that affect the steady-state concentration of active complexes: ABA, ABC, A3B or ABCD. Below we use $k_{sig}$ to represent these processes are we are particularly interested in how signaling time-scales affect the validity of our equilibrium models. Assuming steady-state conditions: i.e. that the rate of assembly and disassembly of the active complex ABA is equal:

$$2k_1[A][B] = \left(k_{sig} + \frac{2(k_{-1})^2}{\alpha k_1} \times \frac{1}{[A]^{surf}}\right)[ABA] \tag{S77}$$

Solving for [B]:

$$[B] = \frac{[ABA]}{[A]}\left(\frac{k_{sig} + \frac{2(k_{-1})^2}{\alpha k_1}\frac{1}{[A]^{surf}}}{2k_1}\right) \tag{S78}$$

At the signaling $EC_{50}$, if we assume 50% maximum complex formation, then the conservation of mass yields concentrations of all free species in equation S78:

$$[ABA]_{EC50} = [A]_t/4 \tag{S79}$$

$$[A]_{EC50} = [A]_t/2 \tag{S80}$$

$$[B]_{EC50} = [B]_t - [A]_t/4 \tag{S81}$$

Replacing all free expressions in equation S78 with equations S79-81 and simplifying we obtain:

$$[B]_t - [A]_t/4 = \frac{[A]_t/4}{[A]_t/2}\left(\frac{k_{sig} + \frac{2(k_{-1})^2}{\alpha k_1}\frac{1}{[A]^{surf}/2}}{2k_1}\right) \tag{S82}$$

Simplifying and solving for $[B]_t$ which gives us our $EC_{50}$

$$EC_{50} = \frac{[A]_t}{4} + \frac{k_{sig}}{4k_{+1}} + \frac{(K_1)^2}{\alpha[A]_t^{surf}} \tag{S83}$$

##### *ii. ABC Complex $EC_{50}$ (Assuming $k_{AB} < k_{BC}$)*

First putting the law of mass action[13] in terms of microscopic rate constants

$$\frac{k_{-AB}k_{-BC}}{\alpha k_{+AB}k_{+BC}}\frac{V_{surf}}{V_{soln}} = \frac{[A][B][C]}{[ABC]} \tag{S84}$$

Defining the formation of AB as the first committed step in the assembly of ABC (this necessitates that the formation of AB must be faster than BC as such for the purposes of this derivation we assume $k_{AB} << k_{BC}$ (in the next section we assume the converse))

$$2k_{AB}[A][B] = \left(k_{sig} + \frac{k_{-AB}k_{-BC}}{\alpha k_{+BC}} \times \frac{1}{[C]^{surf}}\right)[ABC] \tag{S85}$$

Solving for [B]

$$[B] = \frac{[ABC]}{[A]} = \left( \frac{k_{sig} + \frac{k_{-AB}k_{-BC}}{\alpha k_{+BC}} \frac{1}{[C]^{surf}}}{k_{+AB}} \right) \quad S86$$

At the signaling $EC_{50}$ all free concentrations are known

$$[ABC]_{EC50} = [A]_t/2 \quad S87$$

$$[A]_{EC50} = [A]_t/2 \quad S88$$

$$[B]_{EC50} = [B]_t - [A]_t/2 \quad S89$$

$$[C]_{EC50} = [C]_t - [A]_t/2 \quad S90$$

Replacing all free concentrations in Mass Conservation equation for $[B]_t$ [13]with equations S87-S90:

$$[B]_t - [A]_t/2 = \frac{[A]_t/2}{[A]_t/2} \left( \frac{k_{sig} + \frac{k_{-AB}k_{-BC}}{\alpha k_{+BC}} \frac{1}{[C]_t^{surf} - [A]_t^{surf}/2}}{k_{+AB}} \right) \quad S91$$

Simplifying and solving for $[B]_t$ which yields our $EC_{50}$:

$$EC_{50} = \frac{[A]_t}{2} + \frac{k_{sig}}{k_{+AB}} + \frac{K_{AB}K_{BC}}{\alpha\left([C]_t^{surf} - [A]_t^{surf}/2\right)} \quad S92$$

##### *iii. ABC Complex $EC_{50}$ (Assuming $k_{BC} > k_{AB}$)*

Defining the formation of BC as the first committed step in the assembly of ABC (this necessitates that the formation of BC must be faster than AB as such for the purposes of this derivation we assume $k_{BC} >> k_{AB}$ (in the previous section we assumed the converse))

$$2k_{BC}[A][B] = \left( k_{sig} + \frac{k_{-AB}k_{-BC}}{\alpha k_{+BC}} \times \frac{1}{[C]^{surf}} \right) [ABC] \quad S93$$

Solving for [B]

$$[B] = \frac{[ABC]}{[A]} \left( \frac{k_{sig} + \frac{k_{-AB}k_{-BC}}{\alpha k_{+BC}} \frac{1}{[C]^{surf}}}{k_{+BC}} \right) \quad S94$$

At the signaling $EC_{50}$ all free concentrations are known

$$[ABC]_{EC50} = [A]_t/2 \quad S95$$

$$[A]_{EC50} = [A]_t/2 \quad \text{S96}$$

$$[B]_{EC50} = [B]_t - [A]_t/2 \quad \text{S97}$$

$$[C]_{EC50} = [C]_t - [A]_t/2 \quad \text{S98}$$

Replacing all free concentrations in Mass Conservation equation for $[B]_t$[13] with equations S95-S98:

$$EC_{50} = \frac{[A]_t}{2} + \frac{[A]_t/2}{[C]_t - [A]_t/2}\frac{k_{sig}}{k_{+BC}} + \frac{K_{AB}K_{BC}}{\alpha\left([C]_t^{surf} - [A]_t^{surf}/2\right)} \quad \text{S99}$$

Interestingly when the limiting reagent (here defined as A) is different from the receptor involved in the rate determining step the "kinetic limit" on potency is no longer $k_{sig}/k_{on}$ and the ratio of receptors contributes. This is an important clarification of muti-valent receptor systems relative to what would be expected from simple extension of Michaelis-Menten "kinetic limits" to ternary systems by analogy.

##### *iv. $A_3B$ Complex $EC_{50}$*

Writing out the law of mass action for $A_3B$ type complexes in terms of microscopic rate constants:

$$\frac{V_{surf}}{V_{soln}}\frac{k_{-1}^3}{\alpha\beta k_1^3} = \frac{[A][B][A][A]}{[A_3B]} \quad \text{S100}$$

Defining the formation of AB as the RDS the steady-state equation can be written as:

$$3k_1[A][B] = \left(k_{sig} + \frac{3(k_{-1})^3}{\alpha\beta(k_1)^2} \times \frac{1}{([A]^{surf})^2}\right)[A_3B] \quad \text{S101}$$

Solving for [B]

$$[B] = \frac{[A_3B]}{[A]}\left(\frac{k_{sig} + \frac{2(k_{-1})^3}{\alpha\beta(k_1)^2}\frac{1}{([A]^{surf})^2}}{3k_1}\right) \quad \text{S102}$$

At the signaling $EC_{50}$ the concentrations of all free species are known:

$$[ABA]_{EC50} = [A]_t/6 \quad \text{S103}$$

$$[A]_{EC50} = [A]_t/2 \quad \text{S104}$$

$$[B]_{EC50} = [B]_t - [A]_t/6 \quad \text{S105}$$

Replacing all free expressions in S102 with S103-S105:

$$[B]_t - [A]_t/6 = \frac{[A]_t/6}{[A]_t/2}\left(\frac{k_{sig} + \frac{2(k_{-1})^3}{\alpha\beta(k_1)^2}\frac{1}{([A]^{surf}/2)^2}}{3k_1}\right) \quad \text{S106}$$

Solving for $[B]_t$ and simplifying we obtain:

$$EC_{50} = \frac{[A]_t}{6} + \frac{k_{sig}}{9k_{+1}} + \frac{4}{3}\frac{(K_1)^2}{\alpha\beta([A]^{surf})^2} \quad \text{S107}$$

##### *v. $A_nB$ Complex $EC_{50}$*

Writing out the law of mass action for $A_3B$ type complexes in terms of microscopic rate constants:

$$\left(\frac{V_{surf}}{V_{soln}}\right)^n \left(\frac{k_{-1}}{k_{+1}}\right)^n = \frac{[A]^n[B]}{[A_nB]} = \frac{"k_{off}"}{"k_{on}"} \qquad \text{S108}$$

Defining the formation of AB as the RDS the steady-state equation is:

formation of AB as the RDS the steady-state equation is:

$$nk_1[A][B] = \left(k_{sig} + \frac{3(k_{-1})^n}{(k_1)^{n-1}} \times \frac{1}{([A]^{surf})^{n-1}}\right)[A_nB] \qquad \text{S109}$$

Solving for [B]

$$[B] = \frac{[A_nB]}{[A]}\left(\frac{k_{sig} + \frac{2(k_{-1})^n}{(k_1)^{n-1}}\frac{1}{([A]^{surf})^{n-1}}}{nk_1}\right) \qquad \text{S110}$$

At the signaling $EC_{50}$ the concentrations of all free species are known:

$$[ABA]_{EC50} = [A]_t/2n \qquad \text{S111}$$

$$[A]_{EC50} = [A]_t/2 \qquad \text{S112}$$

$$[B]_{EC50} = [B]_t - [A]_t/2n \qquad \text{S113}$$

Replacing all free expressions in S110 with equations S111-S113:

$$[B]_t - [A]_t/2n = \frac{[A]_t/2n}{[A]_t/2n}\left(\frac{k_{sig} + \frac{2(k_{-1})^n}{(k_1)^{n-1}}\frac{1}{\left([A]_t^{surf}/2\right)^{n-1}}}{nk_1}\right) \qquad \text{S114}$$

Solving for $[B]_t$ and simplifying we obtain:

$$EC_{50} = \frac{[A]_t}{2n} + \frac{k_{sig}}{n^2k_{+1}} + K_1\frac{1}{n}\left(\frac{2K_1}{[A]_t^{surf}}\right)^{n-1} \qquad \text{S115}$$

***vi-1. ABCD Complex $EC_{50}$ (Assuming $k_{AB}$ > $k_{BC}$ or $k_{BD}$)***

Writing out the law of mass action in terms of individual rate constants:

$$\frac{k_{-AB}k_{-BC}k_{-BD}}{\alpha\beta\gamma\delta k_{+AB}k_{+BC}k_{+BD}}\left(\frac{V_{surf}}{V_{soln}}\right)^2 = \frac{[A][B][C][D]}{[ABCD]} \quad \text{S116}$$

Defining the formation of AB as the first committed step in the assembly of ABCD (this necessitates that the formation of AB must be faster than BC and BD as such for the purposes of this derivation we assume $k_{AB}$ >> $k_{BC}$,$k_{BD}$ (in the next section we assume the converse))

$$k_{AB}[A][B] = \left(k_{sig} + \frac{k_{-AB}k_{-BC}k_{-BD}}{\alpha\beta\gamma\delta k_{+BC}k_{+BD}} \times \frac{1}{[C]^{surf}[D]^{surf}}\right)[ABCD] \quad \text{S117}$$

Solving for [B]

$$[B] = \frac{[ABCD]}{[A]}\left(\frac{k_{sig} + \frac{k_{-AB}k_{-BC}k_{-BD}}{\alpha\beta\gamma\delta k_{+BC}k_{+BD}}\frac{1}{[C]^{surf}[D]^{surf}}}{k_{+AB}}\right) \quad \text{S118}$$

At the signaling $EC_{50}$ all the free concentrations are known

$$[ABCD]_{EC50} = [A]_t/2 \quad \text{S119}$$

$$[A]_{EC50} = [A]_t/2 \quad \text{S120}$$

$$[B]_{EC50} = [B]_t - [A]_t/2 \quad \text{S121}$$

$$[C]_{EC50} = [C]_t - [A]_t/2 \quad \text{S122}$$

$$[D]_{EC50} = [D]_t - [A]_t/2 \quad \text{S123}$$

Replacing S118 for the free concentrations in S119-S123:

$$[B]_t - [A]_t/2 = \frac{[A]_t/2}{[A]_t/2}\left(\frac{k_{sig} + \frac{k_{-AB}k_{-BC}k_{-BD}}{\alpha\beta\gamma\delta k_{+BC}k_{+BD}}\frac{1}{\left([C]_t^{surf} - [A]_t^{surf}/2\right)\left([D]_t^{surf} - [A]_t^{surf}/2\right)}}{k_{+AB}}\right) \quad \text{S124}$$

Simplifying and Solving for $[B]_t$ which yields the $EC_{50}$

$$EC_{50} = \frac{[A]_t}{2} + \frac{k_{sig}}{k_{+AB}} + \frac{K_{AB}K_{BC}K_{BD}}{\alpha\beta\gamma\delta\left([C]_t^{surf} - [A]_t^{surf}/2\right)\left([C]_t^{surf} - [A]_t^{surf}/2\right)} \quad \text{S125}$$

***vi-2. ABCD Complex $EC_{50}$ (Assuming $k_{AB}$ < $k_{BC}$ or $k_{BD}$)***

Defining the formation of BD as the first committed step in the assembly of ABCD (this necessitates that the formation of BD must be faster than AB and BC as such for the purposes of this derivation we assume $k_{BD} >> k_{AB},k_{BC}$ (this is equivalent to $k_{BC} >> k_{AB},k_{BD}$ and in the previous section we assume the converse))

$$k_{BD}[B][D] = \left(k_{sig} + \frac{k_{-AB}k_{-BC}k_{-BD}}{\alpha\beta\gamma\delta k_{+AB}k_{+BC}} \times \frac{1}{[A]^{surf}[C]^{surf}}\right)[ABCD] \qquad \text{S126}$$

Solving for [B]

$$[B] = \frac{[ABCD]}{[D]}\left(\frac{k_{sig} + \frac{k_{-AB}k_{-BC}k_{-BD}}{\alpha\beta\gamma\delta k_{+AB}k_{+BC}} \times \frac{1}{[A]^{surf}[C]^{surf}}}{k_{+BD}}\right) \qquad \text{S127}$$

At the signaling $EC_{50}$ all the free concentrations are known

$$[ABCD]_{EC50} = [A]_t/2 \qquad \text{S128}$$

$$[A]_{EC50} = [A]_t/2 \qquad \text{S129}$$

$$[B]_{EC50} = [B]_t - [A]_t/2 \qquad \text{S130}$$

$$[C]_{EC50} = [C]_t - [A]_t/2 \qquad \text{S131}$$

$$[D]_{EC50} = [D]_t - [A]_t/2 \qquad \text{S132}$$

Replacing equations S128-S132 for the free concentrations in S127 we obtain:

$$[B]_t - [A]_t/2 = \frac{[A]_t/2}{[D]_t - [A]_t/2}\left(\frac{k_{sig} + \frac{k_{-AB}k_{-BC}k_{-BD}}{\alpha\beta\gamma\delta k_{+AB}k_{+BC}} \frac{1}{\left([A]_t^{surf}/2\right)\left([C]_t^{surf} - [A]_t^{surf}/2\right)}}{k_{+BD}}\right) \qquad \text{S133}$$

Simplifying and Solving for $[B]_t$ which yields the $EC_{50}$

$$EC_{50} = \frac{[A]_t}{2} + \frac{[A]_t}{2[D]_t - [A]_t}\frac{k_{sig}}{k_{+BD}} + \frac{K_{AB}K_{BC}K_{BD}}{\alpha\beta\gamma\delta\left([C]_t^{surf} - [A]_t^{surf}/2\right)\left([D]_t^{surf} - [A]_t^{surf}/2\right)} \qquad \text{S134}$$

| *Table 1. Summary of $EC_{50}$ Including a Kinetic Limit (Steady-State Assumption)* | |
|---|---|
| MODEL | $EC_{50}$ |
| [ABA] | $EC_{50} = \frac{[A]_t}{4} + \frac{k_{sig}}{4k_{+1}} + \frac{(K_1)^2}{\alpha [A]_t^{surf}}$ |
| [ABC] | $EC_{50} = \frac{[A]_t}{2} + \frac{k_{sig}}{k_{+AB}} + \frac{K_{AB}K_{BC}}{\alpha\left([C]_t^{surf} - [A]_t^{surf}/2\right)}$ <br> or <br> $EC_{50} = \frac{[A]_t}{2} + \frac{k_{sig}}{k_{+BC}} + \frac{K_{AB}K_{BC}}{\alpha\left([C]_t^{surf} - [A]_t^{surf}/2\right)}$ |
| $[A_3B]$ | $EC_{50} = \frac{[A]_t}{6} + \frac{k_{sig}}{9k_{+1}} + \frac{4}{3}\frac{(K_1)^2}{\alpha\beta([A]^{surf})^2}$ |
| $[A_nB]$ | $EC_{50} = \frac{[A]_t}{2n} + \frac{k_{sig}}{n^2 k_{+1}} + K_1 \frac{1}{n}\left(\frac{2K_1}{[A]_t^{surf}}\right)^{n-1}$ |
| [ABCD] | $EC_{50} = \frac{[A]_t}{2} + \frac{k_{sig}}{k_{+AB}} + \frac{K_{AB}K_{BC}K_{BD}}{\alpha\beta\gamma\delta\left([C]_t^{surf} - [A]_t^{surf}/2\right)\left([C]_t^{surf} - [A]_t^{surf}/2\right)}$ <br> or <br> $EC_{50} = \frac{[A]_t}{2} + \frac{[A]_t}{2[C]_t - [A]_t}\frac{k_{sig}}{k_{+BC}} + \frac{K_{AB}K_{BC}K_{BD}}{\alpha\beta\gamma\delta\left([C]_t^{surf} - [A]_t^{surf}/2\right)\left([D]_t^{surf} - [A]_t^{surf}/2\right)}$ <br> or <br> $EC_{50} = \frac{[A]_t}{2} + \frac{[A]_t}{2[D]_t - [A]_t}\frac{k_{sig}}{k_{+BD}} + \frac{K_{AB}K_{BC}K_{BD}}{\alpha\beta\gamma\delta\left([C]_t^{surf} - [A]_t^{surf}/2\right)\left([D]_t^{surf} - [A]_t^{surf}/2\right)}$ |
